# Cortical Origin of Theta Error Signals

**DOI:** 10.1101/2023.06.27.546752

**Authors:** Beatriz Herrera, Amirsaman Sajad, Steven P. Errington, Jeffrey D. Schall, Jorge J. Riera

**Author notes:** Corresponding Authors: Jorge J. Riera.

## Abstract

A multi-scale approach elucidated the origin of the error-related-negativity (ERN), with its associated theta-rhythm, and the post-error-positivity (Pe) in macaque supplementary eye field (SEF). Using biophysical modeling, synaptic inputs to layer-3 (L3) and layer-5 (L5) pyramidal cells (PCs) were optimized to account for error-related modulation and inter-spike intervals. The intrinsic dynamics of dendrites in L5 but not L3 PCs generate theta rhythmicity with random phase. Saccades synchronized the phase of this theta-rhythm, which was magnified on errors. Contributions from L5 PCs to the laminar current source density (CSD) observed in SEF were negligible. The CSD derived from L3 PCs could not explain the observed association between their error-related spiking modulation and scalp-EEG. Laminar CSD comprises multipolar components, with dipoles explaining ERN features, and quadrupoles reproducing those for Pe. The presence of monopoles indicates diffuse activation. These results provide the most advanced explanation of the cellular mechanisms generating the ERN.

## Introduction

Cognitive control involves the suppression of automatic or impulsive actions and error monitoring for successful goal-directed behavior. Disorders such as attention-deficit hyperactivity disorder (ADHD)(Armstrong and Munoz 2003; Hanisch et al. 2006), obsessive-compulsive disorder (OCD) (Penadés et al. 2007), and schizophrenia (Donohoe et al. 2006), involve insufficient cognitive control (Aron et al. 2003). Human and macaque electrophysiological studies have characterized the scalp potentials associated with error monitoring (Gehring et al. 2012), the error-related negativity (ERN) associated with prominent midfrontal theta oscillations (Cavanagh and Frank 2014; Cohen 2014). Although the ERN is known to originate from medial frontal areas (Stuphorn et al. 2000; Garavan et al. 2003; Ito et al. 2003; Emeric et al. 2008, 2010; Gehring et al. 2012; Scangos et al. 2013; Sajad et al. 2019; Fu et al. 2023), the cellular-level mechanisms producing these signals and their involvement in midfrontal theta generation are unknown. A better understanding of the mechanisms of error monitoring at the microcircuit level will provide more insights into the underlying intricacies of neurological disorders and hence aid their diagnosis and treatment by mechanistically defining ERN biomarkers.

Performance evaluation indexed by the ERN can be investigated with the stop-signal task (Verbruggen and Logan 2009). Specific neurons in the supplementary eye field (SEF) signal gaze errors, causing an imprint in the local field potential (LFP) (Stuphorn et al. 2000; Emeric et al. 2010). Recently, we have used linear electrode arrays to characterize the laminar organization of neural processing in SEF (Sajad et al. 2019, 2022). We found that most error-related neurons have broad spikes, consistent with pyramidal cells (PCs), and that the variability in spiking of neurons in layers 2 and 3, but not in layers 5 and 6, is statistically associated with the variability of the ERN. It remains unclear whether these error-related PCs contribute directly to the LFP in SEF or cause large circuital activation that are then visible in LFP. What types of brain source components in SEF generate the ERN, and potentially Pe, is another unsolved question. Also, previous studies have suggested a microcircuit origin for the theta oscillation in midfrontal cortical areas (Cohen 2014), involving positive feedback between L3 and L5 PCs as well as an inhibitory close loop by Martinotti cells. Mechanisms linking error-related PCs to theta oscillation are still elusive. Current source density (CSD) and time-frequency analysis methods offer insights into layer-specific contributions but cannot resolve the distinct neuronal populations. In our opinion, biophysically detailed modeling of the activity of L3 and L5 PCs in SEF, combined with mesoscopic brain source models, is required to resolve such cell-specific mechanisms from multiscale electrophysiological data.

Here, we combined detailed biophysical modeling of individual PCs with neural data recorded in SEF from two macaque monkeys performing the stop-signal task (Godlove et al. 2014; Sajad et al. 2019). The spatio-temporal pattern of excitatory (NMDA and AMPA) pre-synaptic inputs were optimized in models of L3 and L5 PCs to replicate observed error-related modulation and inter-spike interval profiles before and during the testing trials. The LFP across cortical layers derived from the parameterized model of L5 but not of L3 PCs produced a significant increase in theta power on error versus correct trials. Although peaking during the ERN, the current density derived from the simulated PCs provided a negligible contribution to both the CSD in SEF and the scalp-ERN. The observed current density included a well-defined dipolar component explaining the ERN and a significant quadrupolar contribution to the Pe component. Overall, these results suggest localized activation in SEF underlying the ERN, but the Pe component might be more diffuse in SEF or perhaps involve other cortical regions. The origin of a large monopolar source found in SEF is yet to be explained. By translating across scales, these findings offer unprecedented insights into the mechanisms of cognitive control and the origin of the ERN.

## Materials and Methods

### Experimental Model and Subject Details

Data were collected from one male bonnet macaque (Eu, Macaca radiata, ∼8.8 kg) and one female rhesus macaque (X, Macaca Mulatta, ∼6.0 kg) performing a saccade countermanding stop-signal task (Hanes and Schall 1995; Godlove et al. 2014). Monkeys were cared for in accordance with the United States Department of Agriculture and Public Health Service Policies on Human Care and Use of Laboratory Animals. All procedures were performed with supervision and approval from the Vanderbilt Institutional Animal Care and Use Committee.

### Animal Care and Surgical Procedures

Anatomic images were acquired with a Philips Intera Achieva 3 Tesla scanner using SENSE Flex-S surface coils placed above and below the head. T1-weighted gradient-echo structural images were obtained with a 3D turbo field echo anatomical sequence (TR = 8.729 ms; 130 slices, 0.70 mm thickness). Anatomical images guided the placement of Cilux recording chambers in the correct area. Chambers were implanted normal to the cortex (Monkey Eu: 17°; 777 Monkey X: 9°; relative to stereotaxic vertical) centered on the midline, 30mm (Monkey Eu) and 778 28mm (Monkey X) anterior to the interaural line. Surgical procedures have been previously described (Godlove, Garr, et al. 2011).

### Saccade Countermanding Stop-Signal Task

Monkeys performed a stop-signal saccade countermanding task (monkeys Eu and X) (Fig. 1A). All trials started with the presentation of a central fixation spot in the form of a square. Monkeys were required to hold fixation for a variable interval after which the center of the square was extinguished. Simultaneously, a peripheral target at either the right or left of the fixation spot was presented. On no-stop-signal trials, monkeys were required to generate a saccade to a peripheral target, whereupon after 600 ± 0 ms, a high-pitched auditory feedback tone was delivered, and 600 ± 0 ms later, fluid reward was provided. On stop-signal trials, following the target presentation and after a variable stop-signal delay (SSD), the fixation spot was re-illuminated instructing the monkey to inhibit the planned saccade. In the trials where the monkey successfully canceled the saccade to the peripheral target, the same high-pitch tone was presented after a 1,500 ± 0 ms hold time followed, after 600 ± 0 ms, by fluid reward. SSD was adjusted such that monkeys successfully canceled the saccade in ∼50% of the trials. Noncanceled errors occurred when monkeys generated a saccade despite the appearance of stop-signal. In these trials, a low-pitch tone was presented 600 ± 0 ms after the saccade and no fluid reward was delivered.

**Fig. 1:**
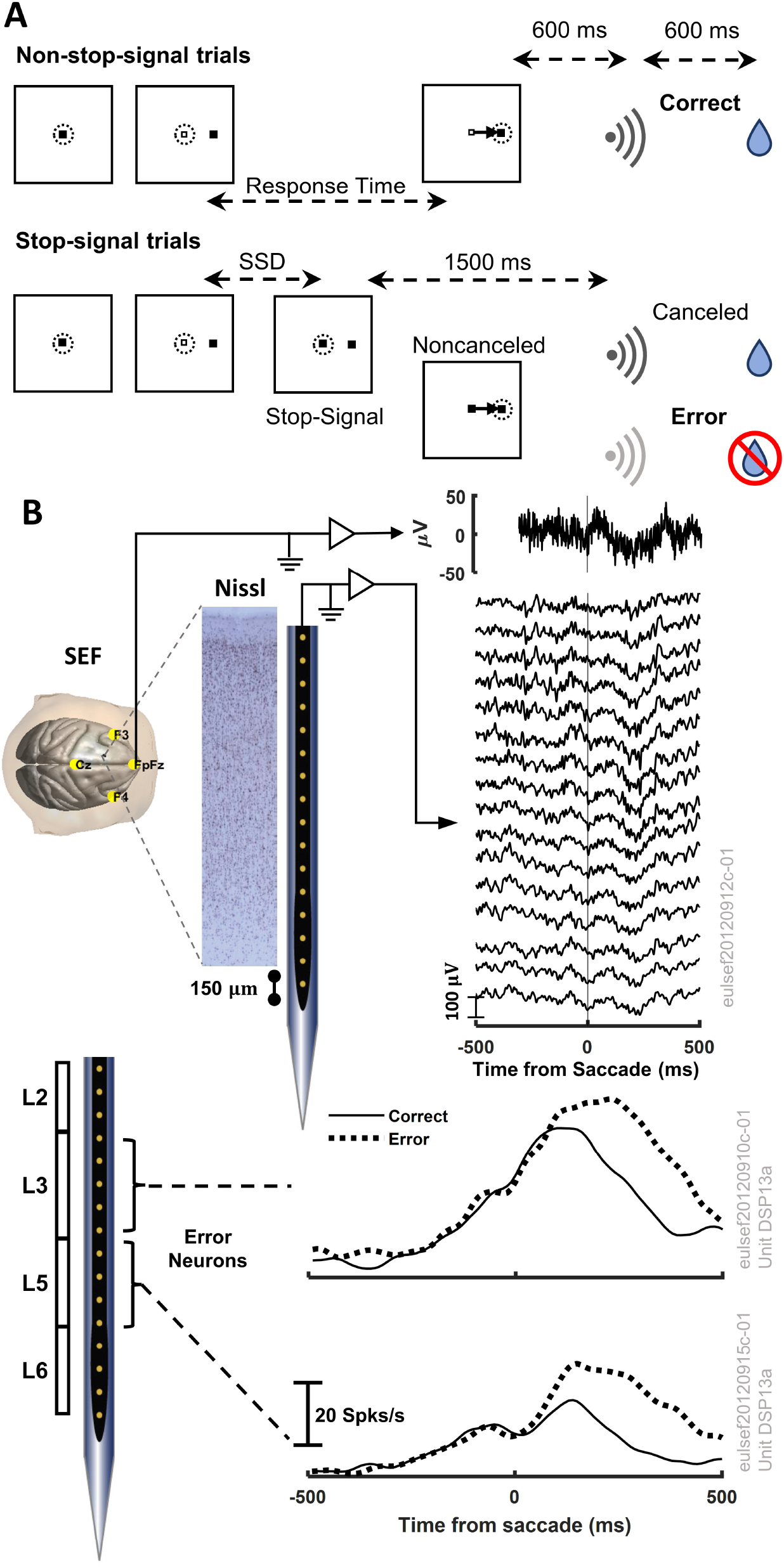
Experimental procedures and methodology. A. Stop-signal saccade countermanding task. All trials started with the presentation of a square fixation marker. Monkeys were required to hold fixation for a variable interval after which the center of the square was extinguished simultaneous with presentation of a peripheral target on the right or left. On no-stop-signal trials, monkeys shifted gaze to the target, whereupon after 600 ± 0 ms a high-pitched tone was delivered followed 600 ± 0 ms later by fluid reward. On stop-signal trials, a variable stop-signal delay (SSD) after target presentation the center of the fixation spot was re-illuminated instructing the monkey to inhibit the planned saccade. If monkeys canceled the saccade, the high-pitch tone was presented after 1,500 ± 0 followed 600 ± 0 ms later by fluid reward. SSD was adjusted such that monkeys successfully canceled the saccade in ∼50% of the trials. if monkeys produced a noncanceled error, a low-pitch tone was presented 600 ± 0 ms after the saccade and no fluid reward was delivered. B. Schematic of concurrent EEG and LFP recording in SEF used to calculate theta power and current source density after saccades (top) and mean spike rate of representative L3 and L5 putative error PCs (bottom).

### Cortical mapping and electrode placement

Chambers implanted over the medial frontal cortex were mapped using tungsten microelectrodes (2-4 MΩ, FHC, Bowdoin, ME) to apply 200ms trains of biphasic micro-stimulation (333 Hz, 200 µs pulse width). The SEF was identified as the area from which saccades could be elicited using < 50 µA of current (Schlag and Schlag-Rey 1987; Schall 1991). In both monkeys, the SEF chamber was placed over the left hemisphere. The dorsomedial location of the SEF makes it readily accessible for linear electrode array recordings across all cortical layers. A total of five penetrations were made into the cortex—two in monkey Eu, and three in monkey X. Three of these penetration locations were perpendicular to the cortex. In monkey Eu, the perpendicular penetrations sampled activity at site P1, located 5 mm lateral to the midline and 31 mm anterior to the interaural line. In monkey X, the perpendicular penetrations sampled activity at sites P2 and P3, located 5 mm lateral to the midline and 29 and 30 mm anterior to the interaural line, respectively. However, during the mapping of the bank of the cortical medial wall, we noted both monkeys had chambers placed ∼1 mm to the right respective to the midline of the brain. This was confirmed through co-registered CT/MRI data. Subsequently, the stereotaxic estimate placed the electrodes at 4 mm lateral to the cortical midline opposed to the skull-based stereotaxic midline.

### Spiking activity and local field potential recordings

During recordings, monkeys sat in enclosed primate chairs with heads restrained 45 cm from a CRT monitor (Dell P1130, background luminance of 0.10 *cd*/*m*^2^, 70 Hz) subtending 46° × 36° of visual angle. Daily recording protocols were consistent across monkeys and sessions. After advancing the electrode array to the desired depth, electrodes were allowed to settle for three to four hours to ensure stable recordings.

Spiking activity and local field potentials (LFPs) were recorded from SEF using a 24-channel U probe (Plexon, Dallas, TX) with 150 *μm* inter-electrode distance. The U probes had 100 mm probe length with 30 mm reinforced tubing, 210 *μm* probe diameter, 30° tip angle, 500 *μm* to first contact. Contacts were referenced to the probe shaft and grounded to the metal headpost. All data were streamed to a data acquisition system (MAP, Plexon, Dallas, TX). Time stamps of trial events were recorded at 500 Hz. Eye position data were streamed to the Plexon computer at 1 kHz using an EyeLink 1000 infrared eye-tracking system (SR Research, Kanata, Ontario, Canada). LFP and spiking data were processed with unity-gain high-input impedance head stages (HST/32o25-36P-TR, Plexon).

LFP data were bandpass filtered at 0.2–300 Hz and amplified 1000 times with a Plexon preamplifier and digitized at 1 kHz. Neuronal spiking data were bandpass filtered between 100 Hz and 8 kHz and amplified 1000 times with a Plexon preamplifier, filtered in software with a 250 Hz high-pass filter, and amplified an additional 32,000 times. Waveforms were digitized from − 200 to 1200 *μs* relative to threshold crossings at 40 kHz. Thresholds were typically set at 3.5 standard deviations from the mean. Single units were sorted online using a software window discriminator and refined offline using principal components analysis implemented in Plexon offline sorter.

### Cortical depth and layer assignment

Depth alignment and laminar assignment were performed across sessions as described in Godlove et al. (2014). Briefly, flashed visual stimulation was delivered to the monkeys between sessions. Recording sessions were aligned relative to the initial visually evoked sink observed on the laminar CSD using an automated depth alignment algorithm. The procedure minimized the differences between the averaged visually evoked CSD across sessions in the 50-100 ms window after the visual stimulus onset. The minimum of the initial visually evoked sink, located in L3, was set as depth zero. Based on this convention, the algorithm identified depths 0.21, 0.36, and 1.02 mm as L1 to L2/3, L3 to L5, and L5 to L6 laminar boundaries, respectively.

### Analysis of spiking activity

Single unit spike rate was estimated on a trial-by-trial basis by calculating the peri-stimulus time histogram (PSTH) of recorded spike trains and convolving them with a Gaussian of zero mean and 10 ms standard deviation. We utilized a bin size of 10 ms to calculate the PSTHs. Trials were defined from -500 ms to 1000 ms relative to saccade initiation time. The average instantaneous spike rate for each recorded unit was obtained by taking the mean across trials.

As previously described by Sajad et al. (2019), error neurons were identified as units showing periods of significant difference between their spiking activity on error and correct trials (referred to as difference function) that exceeded 2 standard deviations above a baseline difference measured during the 300 ms period before target onset and persisted for at least 100 ms, or for 50 ms if the difference exceeded 6 standard deviations above the baseline. Only saccades from error and correct trials with similar reaction time (RT) (within 10 ms) and direction were used for comparison. We excluded from the analysis all error trials in which the stop-signal appeared after saccade initiation time. Trials with unstable spiking activity were also excluded from the analyses.

We study the spiking profiles of error neurons using their ISI distribution in the pre-target and post-saccade periods. ISI distributions were calculated using the function ft_spike_isi() from the FieldTrip toolbox (Oostenveld et al. 2011) with a bin size of 2 ms. For calculating the pre-target ISI distributions, we considered all spikes fired before the target presentation until the beginning of the trial. We calculated the pre-target ISI distribution for error and correct trials individually but found no differences. Thus, we combined all trial types for calculating the final pre-target ISI distribution reported in Fig. 2E and Fig. 3E. We obtained the post-saccade ISI distributions considering all spikes fired after saccade initiation and before the delivery of the feedback tone. ISI distributions were normalized by the total number of trials before calculating the averaged ISI distribution across neurons.

**Fig. 2:**
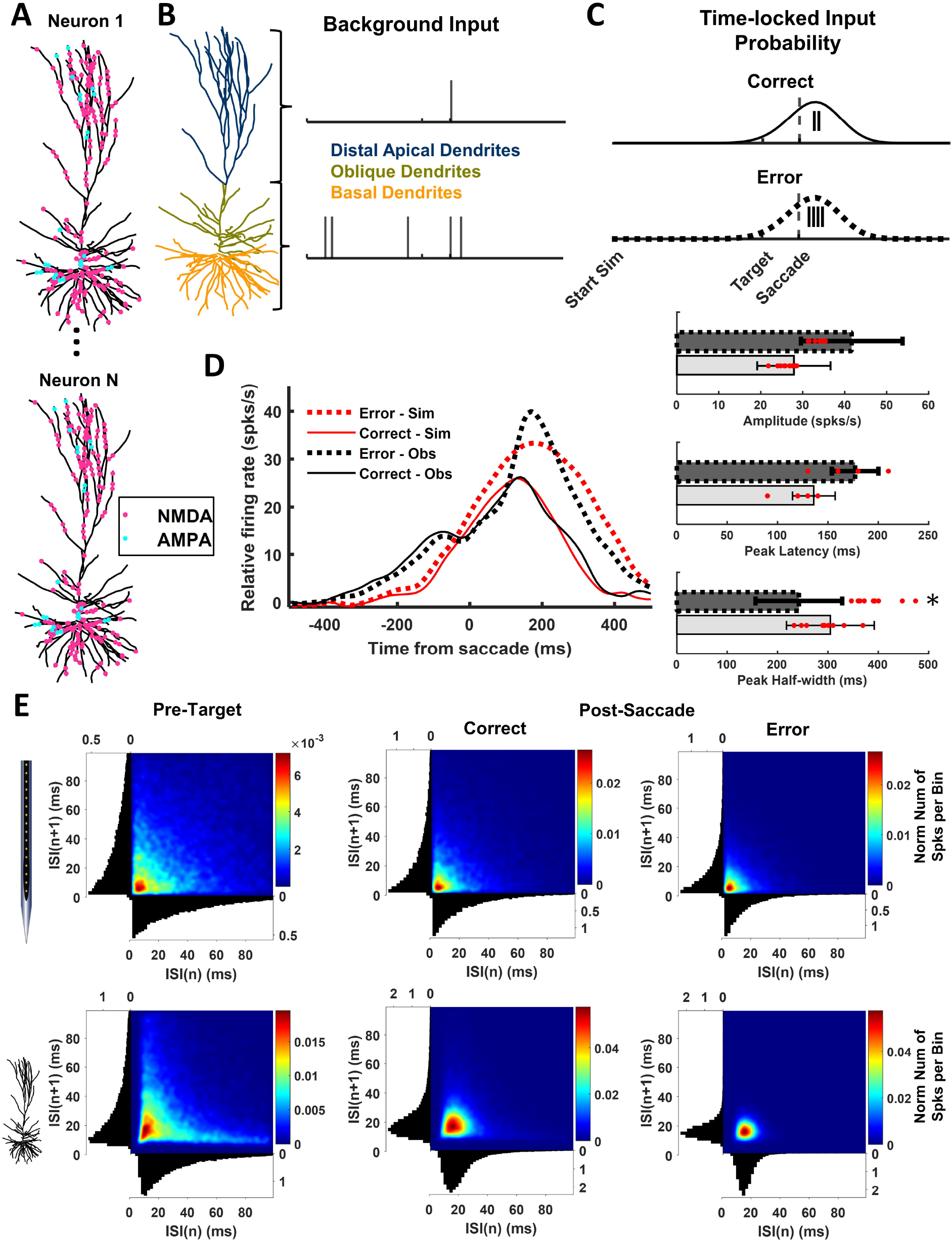
Simulation of L3 error pyramidal cells optimized to replicate observed discharge rates and inter-spike intervals. **A.** Representative randomized locations of NMDA (pink) and AMPA (cyan) synapses on simulated L3 pyramidal cell (ModelDB, accession #238347, 2013_03_06_cell03_789_H41_03, active model cell0603_08_model_602). **B.** Observed baseline spiking statistics were replicated by activating NMDA and AMPA synapses located on the distal apical (dark blue), basal (orange) and oblique (light green) dendrites. The timing of pre-synaptic inputs was drawn from Poisson distributions with a mean of 2 for basal and oblique dendritic synapses and mean of 3.5 for distal apical synapses. **C.** Spiking statistics after saccade initiation were simulated by activating distal apical and basal synapses with spike times drawn from a left skewed normal probability distribution (skewness = -1). To replicate observed post-saccadic error-related modulation, for correct trials 2 spikes were drawn from a distribution with mean 216.6 ms ± standard deviation 141.6 ms, and for error trials 4 spikes, from a distribution with 298.6 ms ± 178.6 ms. The vertical lines indicate the total number of pre-synaptic spikes that each synapse will receive under its associated probability distribution. **D.** Observed (black) and simulated (red) mean spike rate for correct (thin solid) and error (thick dotted) trials (left) with comparisons of observed (black) and simulated (red) peak amplitude, peak latency, and peak half width (right). Based on non-parametric permutation tests the simulated values were not different from observed amplitude (correct trials, p = 0.5107; error, p = 0.0654), peak latency (correct, p = 0.2449; error, p = 0.5449), and peak half width for correct (p = 0.1083) but not error trials (p = 0.00036). **E.** Observed (top) and simulated (bottom) ISI_n+1_ versus ISI_n_ with heatmap indicating the normalized number of spikes count per bin and marginal distributions before target presentation (left) and after correct (middle) and error (right) saccades. Simulated ISI produced the observed bursting pattern of successive ISI.

**Fig. 3:**
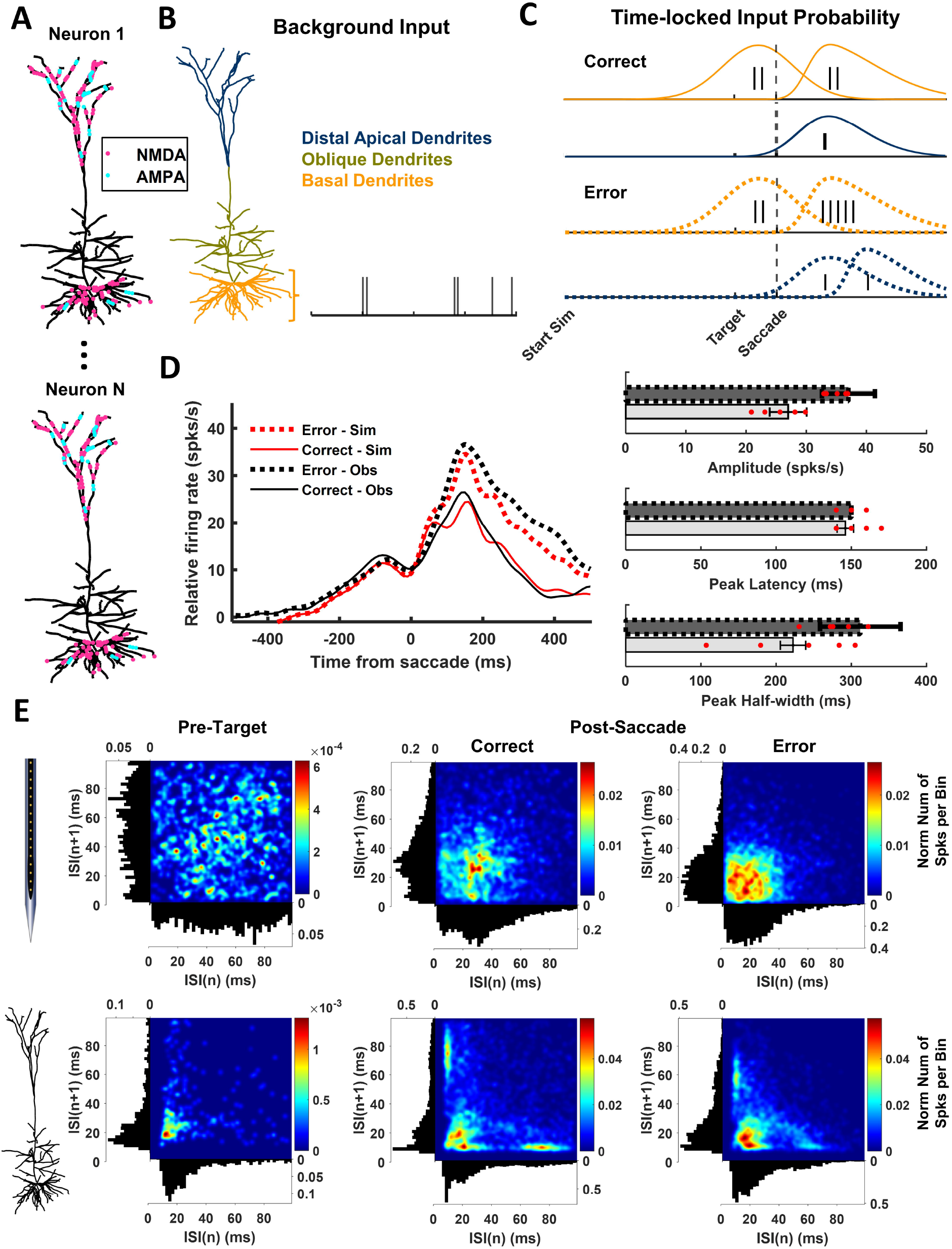
Simulation of L5 error pyramidal cells optimized to replicate observed discharge rates and inter-spike intervals. Conventions as in Fig. 2. **A.** Representative randomized locations of NMDA (pink) and AMPA (cyan) synapses on simulated L5 pyramidal cell (ModelDB, accession #139653, “cell #1”). **B.** Observed baseline spiking statistics were replicated by activating NMDA and AMPA synapses on the basal dendrites with input times drawn from Poisson distributions with a mean of 2. **C.** Spiking statistics before all saccades were simulated by activating basal dendrites with inputs sufficient to produce 2 spikes drawn from a normal distribution (σ = 140 ms) centered 70 ms before saccade initiation. Spiking statistics after correct saccades were simulated with inputs to distal apical dendrites drawn from a right skewed normal distribution (skewness = 2, σ = 200 ms) centered 100 ms after the saccade plus a basal dendrite input drawn from a right skewed normal distribution (skewness = 5, σ = 250 ms) centered 120 ms after the saccade. Spiking statistics after error saccades were simulated by distal apical inputs with the same probability distribution as in correct trials plus a basal input at 120 ms with the same probability distribution as in correct trials sufficient to yield 5 pre-synaptic spikes and a second distal apical input drawn from a right skewed normal distribution (skewness = 5, σ = 250 ms) centered 280 ms after the saccade. **D.** Simulated values were not different from observed amplitude (correct trials, p = 0.4841; error, p = 0.4188), peak latency (correct, p = 0.3783; error, p = 1.0000), and peak half width (correct, p = 0.1553; error, p = 0.3669). **E.** Simulated ISI produced the observed shorter ISI during error trials.

We used a combination of custom-written MATLAB functions (MATLAB 2021b, MathWorks) and the FieldTrip toolbox for analyzing the analyses (Oostenveld et al. 2011).

### Analysis of local field potentials

All analyses were done in MATLAB using custom-written scripts and the FieldTrip toolbox (Oostenveld et al. 2011).

#### ERP and CSD analysis

LFPs were epoched from -500ms to 1,000ms relative to the saccade initiation time, and low-pass filtered at 100 Hz using a two-pass fourth-order Butterworth filter. Recorded trials were separated into correct no-stop-signal and error non-canceled trials. ERPs were time-locked to saccade initiation and baseline corrected to the 200 ms interval preceding the target onset (Godlove et al. 2014).

We computed the CSD from ERPs using the spline-iCSD method (Pettersen et al. 2006) as implemented in the CSDplotter toolbox (https://github.com/espenhgn/CSDplotter) with custom MATLAB (R2021b, The MathWorks) scripts (Herrera et al. 2022).

#### Frequency domain analysis

Time-varying laminar power maps per frequency band were calculated from the LFPs using the Hilbert transform. First, we bandpass filtered the raw LFPs before selecting the trials between 4-8 Hz (θ band), 9-14 Hz (alpha band), 15-29 Hz (beta band), and 30-80 Hz (gamma band), respectively. We constructed 4 Equiripple Bandpass FIR filters using Parks-McClellan optimal FIR filter design, as implemented in firpm() function from MATLAB’s Signal Processing Toolbox 6.14. The optimal filter orders were determined using firpmord() function. Supplemental Fig. S1A shows the magnitude response function of the designed filters. Second, we epoched the filtered LFPs from -500ms to 1,000ms relative to the saccade initiation time. Third, we calculated the Hilbert transform of the filtered LFPs per electrode contact for each trial. Next, we extracted the time-varying power estimates per electrode contact by taking the squared magnitude of the Hilbert transform of the filtered LFPs and then, baseline corrected them to the mean power in the 200 ms interval preceding the target onset. The final time-varying laminar LFP power maps were obtained by taking the mean across the single trial laminar power estimates. Supplemental Fig. S1A illustrates the filtering and epoching procedures and Supplemental Fig. S1B the trial-level processing steps followed to calculate the single trial laminar LFP power estimates.

### Biophysical modeling

#### Pyramidal Cell Models

We simulated the spiking activity of L3 PCs using the previously described model by Eyal et al. (2018) (ModelDB, accession #238347, 2013_03_06_cell03_789_H41_03, active model cell0603_08_model_602). L5 PCs were modeled as previously described in Hay et al. (2011) (ModelDB, accession #139653, “cell #1”), incorporating the modifications of voltage-gated calcium channel densities as in Shai et al. (2015) and Ih channel density distribution as in Labarrera et al. (2018) (Leleo and Segev 2021). Using this modified version of Hay et al. (Leleo and Segev 2021) model allowed us to decrease the bursting activity of the neuron and obtain ISI distributions closer to those observed in the experimental data.

#### Synaptic Inputs

For all simulations, unless otherwise specified, we considered modeled neurons that received excitatory NMDA and AMPA synaptic inputs randomly distributed along their dendrites in clusters of 20 synapses within 20*μm* (Yadav et al. 2012; Kastellakis et al. 2015). The location of the synapses varied for each simulated neuron and trial. The number of NMDA and AMPA synapses for each neuron type was set based on the approximate density of NMDA and AMPA receptors in SEF (area F7d) of macaque monkeys reported in the literature (Geyer et al. 1998; Rapan et al. 2021). We considered the total number of NMDA synapses in the oblique and basal dendrites of L5 PCs (890 synapses) as a reference and determined the number of NMDA synapses in the distal apical dendrites of these neurons based on the relative density of NMDA receptors across the neuron. The total number of AMPA synapses across a simulated L5 PC was calculated based on the ratio of NMDA and AMPA synapses (AMPA-NMDA-ratio: 0.1045) (Rapan et al. 2021). Similarly, we estimated the number of AMPA and NMDA synapses on simulated L3 PCs relative to the set number of NMDA synapses on simulated L5 PCs considering the ratio of L3 to L5-distal-apical AMPA (0.59) (Datta et al. 2015). In summary, we considered a total of 1080, 600, and 1200 NMDA synapses along the basal, oblique, and distal apical dendrites of L3 PCs, and 100, 60, and 120 AMPA synapses, respectively. For L5 PCs, we considered 580 and 444 basal and distal apical dendritic NMDA synapses, and 60 and 132 AMPA synapses, respectively.

*AMPA-*based synaptic currents were modeled as (Mäki-Marttunen et al. 2018): *I_AMPA_* = *w_AMPA_g_AMPA_* (*t*)(*E_AMPA_* – *V*); with *g_AMPA_*(*t*) = (*B_AMPA_* - *A_AMPA_*), and *A_AMPA_* = 0mV. *NMDA-based* synaptic currents were modeled according to the standard formalism(Jahr and Stevens 1990): *I_NMDA_* = *g_NMDA_* (*t*)*w_NMDA_*(*V* – *I_NMDA_*), *g_NMDA_* (*t*) = (*B_NMDA_* - *A_NMDA_*)*f_mg_*(V); *E_NMDA_* = *E_AMPA_* = 0mV with *fmg* (*V*) = 1/(1+0.264 exp (-0.062V)) representing the voltage-dependent magnesium (Mg) block. The equations for *A_i_* and *B_i_* are given by (Mäki-Marttunen et al. 2018): 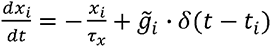 with *x* = {A,B} and i = {*AMPA*, *NMDA*}. *τ_A,AMPA_* = 0.2 ms and *τ_B,AMPA_* = 1.7 ms (Mäki-Marttunen et al. 2018), and *τ_A,NMDA_* = 2 ms and *τ_B,NMDA_* = 100 ms(Jahr and Stevens 1990). For each synapse model, *V* represents the post-synaptic membrane potential, and *t_i_* the onset time of the presynaptic spike. *E_i_*, *g_i_* and *g̃_i_* are the synaptic reversal potential, the gating variable representing the proportion of open channels, and the maximum synaptic conductance, respectively.

#### Estimation of Synaptic Inputs Activation Profiles from Observed Data

We simulated background excitatory inputs coming to L3 and L5 PCs by randomly activating the excitatory synapses on the PC models following a Poisson distribution with a fixed mean. We manually varied the mean of the Poisson process to replicate the observed ISI distribution and spiking activity of recorded L3 and L5 putative error PCs during the pre-target period. Because recorded error neurons from these layers showed large variability in their ISI interval distributions, we did not consider neurons with outlier ISI distributions when obtaining the averaged ISI distribution and spike rate relative to saccade onset. We considered 10 L3 and 5 L5 putative error PCs in the final analyses to constrain the neuronal models.

We modeled time-locked saccade-related inputs as spike generators with a predefined temporal profile relative to the saccade onset time. On a trial-by-trial basis, presynaptic spike times were chosen from a skew-normal distribution (Jones et al. 2007). The number of pre-synaptic temporal profiles and their location along the neuron, as well as their skewness, mean, and standard deviation, were estimated to reproduce the spiking activity and ISI distribution of recorded L3 and L5 putative error PCs relative to saccade onset.

To account for the variable target times and RT observed in the experimental data, we calculated the distribution of target times and saccade times from the experimental data and used these distributions to randomly generate target and saccade onset times for each simulated trial.

#### Analysis of simulated spiking activity

Simulated ISI distributions and spike rates were calculated following the same methodology as for the experimental data. To mimic some of the variability observed in the recorded neurons, we simulated the same number of selected putative L3 (N=10) and L5 (N=5) error PCs and a total of 106 trials, the mean number of trials across sessions in the experimental recordings. In each of these simulations, we randomly varied the location of the pre-synaptic inputs on the modeled neurons, while keeping constant the total number of NMDA and AMPA synapses. Spike times were obtained from the simulated somatic membrane potentials using the peak_detection() function of the Elephant Python package (Denker et al. 2018) with a threshold of 0 mV. To calculate the post-saccade ISI distributions of simulated neurons, we randomly generated the delivery time of the feedback tone for each simulated trial from the experimental distribution of tone times. As for the experimental data, we excluded all spikes fired after the tone.

#### Analysis of simulated field potentials

To study the contribution of L3 and L5 error PCs, we simulated the extracellular field potentials evoked by a population of 625 L3 and 1,000 L5 error PCs under the estimated synaptic inputs. We chose the number of neurons considering the ratio of L3 and L5 PCs in SEF (area F7, ∼>42,500 neurons per mm^3^) (Beul and Hilgetag 2019), the approximate proportion of error neurons in SEF based on all recorded neurons (18 and 16 percent of the neurons recorded from these layers were error neurons, and ∼90% were putative PCs) and the computational costs. We decided to simulate a maximum of 1,000 neurons and scale the magnitude of the laminar CSD obtained from the simulated LFPs. We estimated that a cylindrical cortical column of 3 mm diameter located in SEF would have at least 18,250 L3 and 29,200 L5 error PCs (Beul and Hilgetag 2019), yielding a ratio of L5-to-L3 error PCs of 1.6. Specifically, we calculated the LFP produced by the activity of the neurons at 16 equally spaced vertically aligned points located at the center of a cylindrical cortical column of 3 mm diameter. As in the experiments, the inter-electrode distance was 150 μm. The soma of the neurons was randomly located within the cylindrical cortical column in their associated cortical layers, with height corresponding to the vertical extent in area SEF of lower L3 (700-1100 μm below the pia matter) and L5 (1125-1750 μm). LFPs were calculated from the transmembrane currents using the *point-source approximation* in LFPy (Lindén et al. 2014; Hagen et al. 2018). The *point-source approximation* assumes that each transmembrane current can be represented as a discrete point in space, the center of each neuronal compartment. Considering the extracellular medium is homogeneous and isotropic with an extracellular conductivity *σ_b_*, the extracellular potential Φ(*z_e_*, *t*) at the electrode *z_e_* can be calculated by

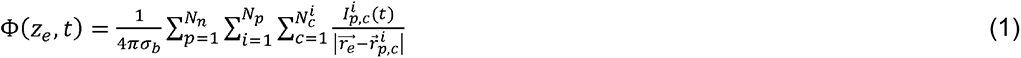

where *N_n_*, *N_p_*, and *N_c_^i^* denote the total number of distinct neuron populations, the number of neurons in the *p-th* population and the number of compartments in the *i-th* neuron of the *p-th* population, respectively. 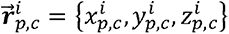 indicates the coordinates of the *c*-*th* compartment of the *i*-*th* neuron in the *p-th* population and 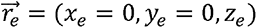 the coordinates of the electrodes. *I_p,c_^i^* (*t*) is the transmembrane current of the *c*-*th* compartment of the *i*-*th* neuron in the *p-th* population.

The LFP was obtained by low pass filtering the extracellular potentials (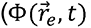) at 100 Hz. LFPs were baseline corrected to the 200 ms interval preceding the target onset. The CSD patterns of the synthetic data sets were calculated using the spline-iCSD method(Pettersen et al. 2006) with the custom MATLAB (R2021b, The MathWorks) scripts used for the experimental data (Herrera et al. 2022). We obtained the time-varying laminar power maps per frequency band from the simulated LFPs using the same analysis pipeline as for the experimental data (Supplementary Fig. S1).

#### Simulations

All biophysical simulations were performed in Python using NEURON 8.0 (Hines et al. 2009) and LFPy 2.2 (Hagen et al. 2018). Data analysis was performed in MATLAB (R2021b, The MathWorks).

### EEG Forward Model

To calculate the EEG potential *V_e_*(***r_e_***, *t*) at the position of the electrodes ***r_e_*** (Fig. 7A), we modeled the monkey’s head as an isotropic and piecewise homogenous volume conductor comprised of the scalp, inner and outer skull, and the cortex surface. For both the experimental and simulated data, we utilized a volume conductor model of the monkey’s head constructed in Brainstorm (Tadel et al. 2011) from the symmetric surfaces provided in the NIMH Macaque Template version 2.0 (Jung et al. 2021) (Fig. 7A). The scalp, skull, and brain conductivities were set as 0.43, 0.0063, and 0.33 S/m (Lee et al. 2015), respectively. In the experimental recordings, only electrodes FpFz, Cz, F3, and F4 were used. Thus, we considered the same electrode positions for our EEG calculations. We obtained the position of the electrodes on the scalp surface of the NIMH Macaque Template using the algorithm from Giacometti et al. (2014) for the EEG 10-10 system.

The EEG potential 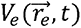 at the electrode position *r⃗_e_* evoked by a continuous field of microscopic electric currents *I*(*r⃗,t*) inside the brain *R* can be calculated by equation (2) (Riera et al. 2012; Herrera et al. 2022):

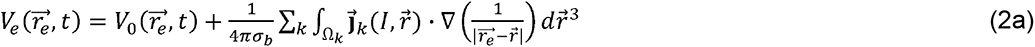

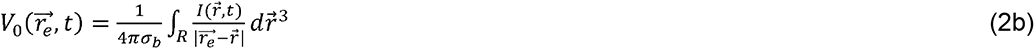

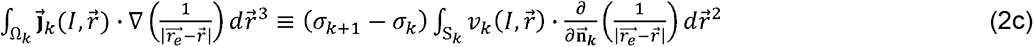

with 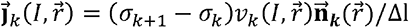 representing the secondary currents defined for each elemental volumetric shell Ω*_k_* (i.e., a surface *S_k_* of thickness Δ*l* → 0). *σ_k_* and *v_k_* (*I*,*r*) denote the conductivity and surface potential of the *k-th* compartment in the head model (i.e., brain (*σ_b_*), skull, and scalp)., and 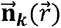 the normal vector to the surface (*S_k_*) of the *k-th* compartment at the location *r⃗*.

Considering that I(*r⃗*,*t*) = *s*(*r*,*t*) for r ∈ *V* and I(*r*,*t*) = 0 otherwise, where V is the volume of the brain region of interest SEF, centered at *r⃗_m_*; and the location of the EEG electrodes (*r⃗_e_*) is far enough from the center *r⃗_m_*, then 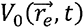 can be calculated as a function of the multipolar moments (Riera et al. 2012). Under the assumption, the EEG forward model can be represented by equation (2a) and the following equation for 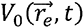 (Riera et al. 2012):

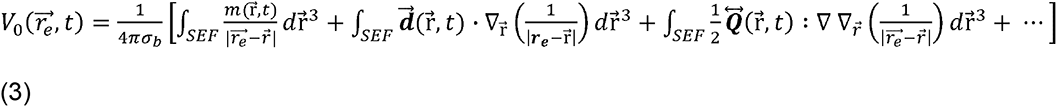

with 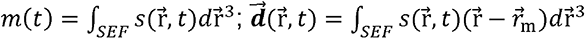; and 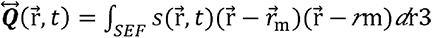.

The first, second, and third terms in equation (3) represent the contribution of the current monopole, dipole and quadrupole, etc., to the EEG, respectively. We demonstrated in Herrera et al. (2022) that the activity of a cortical column can be accurately represented by a single equivalent dipole at the center of the column, whose orientation corresponds to that of the cortical surface and whose temporal dynamic is obtained from the laminar CSD. Using this approach, we calculated the first three current multipole moments from the experimental and simulated CSDs using the following equations:

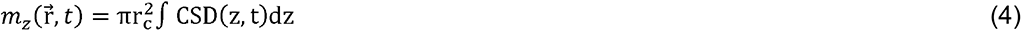

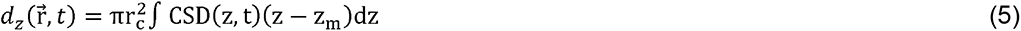

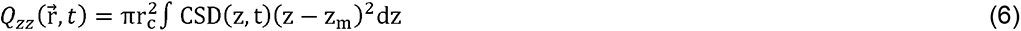

Because of the conservation of current in the neural tissue, the current monopole contribution (equation (4)) should be zero (Nunez and Srinivasan 2009). However, if the CSD is unbalanced, this might be different from zero. Thus, we imposed a monopole and quadrupole moment at the center of the column, at the same location as the equivalent current dipole, to compensate for the current imbalance. This was the case for the observed CSD that had no-zero monopole contribution. The simulated CSD had zero monopole contribution, as expected since the transmembrane currents of compartment models sum to zero at all times. Estimated EEGs were calculated considering there were two symmetric brain sources in SEF, one in each hemisphere. For computing the EEG dipolar contribution, we assumed the orientation of the dipoles corresponded to that of the cortical surface at the dipoles’ location.

### Quantification and Statistical Analysis

#### Spiking activity

We used non-parametric permutation tests for comparing the spike rate features (peak amplitude, peak latency, and peak half-width) between observed and simulated neurons. We used a two-tailed paired t-test to calculate the permutation test statistic and the Monte Carlo method (100,000 permutations for L3 neurons and all possible (40,320) permutations for L5 neurons) for calculating the significance probability, an estimate of the p-value under the permutation distribution. The p-values were reported in the main text or figure captions. The amplitude, latency, and half-width of the peak in the spike rates after the saccade were calculated using the findpeaks() function from MATLAB’s Signal Processing Toolbox.

#### Laminar time-varying field potential frequency power maps

We compared the averaged time-varying laminar power maps of error versus correct trials across sessions (16 sessions, Eu: 6 and X:10) for each frequency band employing nonparametric clustered-based permutation tests (Maris and Oostenveld 2007). We used a two-tailed paired t-test to contrast error versus correct trial averages at the sample level (channel-time-pair samples). Un-smooth power maps were used for the statistical tests. All pairs with t-statistics larger than the critical threshold (*α* = 0.05) were clustered in connected sets based on spatial and temporal adjacency. The cluster-level statistic was calculated by taking the sum of the sample-specific t-statistics within each cluster, and the permutation test statistic was defined as the maximum of the cluster-level test statistic. We utilized the Monte Carlo method for calculating the significance probability, an estimate of the p-value under the permutation distribution. We considered the maximum number of unique permutations for comparison across sessions from each monkey. Significant clusters were determined by comparing their Monte Carlo p-value to an overall two-tailed critical threshold, *α* = 0.01 (0.005 for each tail).

For comparing the simulated error and correct trials, we also employed a nonparametric permutation test but considered a two-tailed unpaired t-test for the sample level statistic (10,000 permutations). In contrast to the experimental data in which we have two experimental conditions per session, only one experimental condition is assigned to each simulated local field potential (between-trial analysis) (Maris and Oostenveld 2007).

## Results

### Electrophysiological recordings

Concurrent scalp EEG and laminar recordings of spiking activity and local field potentials (LFPs) were obtained in supplementary eye field (SEF) of two monkeys (Godlove et al. 2014; Ninomiya et al. 2015) performing the saccade countermanding stop-signal task (Hanes and Schall 1995) (Fig. 1). Briefly, monkeys were required to generate a saccade to a peripheral target, but to inhibit this planned saccade when a stop-signal appeared. Errors occurred when monkeys generated a saccade despite the appearance of the stop-signal. Monkeys produced response errors similar to human participants with homologous ERN features (Godlove, Emeric, et al. 2011; Reinhart et al. 2012). A total of 16 perpendicular sessions were (Eu: 6, X: 10) recorded across monkeys, and resulted in a total of 2,386 trials (Monkey Eu: 1,608; Monkey X: 778) after response-time matching (Godlove et al. 2014; Ninomiya et al. 2015; Sajad et al. 2019). From these sessions, we isolated a total of 293 single units (Eu: 104, X: 189), of which 42 neurons (Eu: 39, X: 3) showed a greater discharge rate following error noncancelled relative to correct saccades (Sajad et al. 2019). The functional properties of these neurons, henceforth referred to as ‘error neurons’, were described previously (Stuphorn et al. 2000; Sajad et al. 2019). Error neurons were divided into putative PCs if their spike waveform had a peak to trough width > 250 μs and interneurons if < 250 μs (Sajad et al. 2019). In total, 37/42 recorded error neurons had broad spike waveforms and were classified as putative PCs. Of these 37, 36 were recorded from L2-L6 of monkey Eu and 1 from L6 of monkey X.

### Reproducing the spiking activity of L3 and L5 error putative pyramidal cells

To evaluate the role of error PCs in the midfrontal theta and ERN generation, we first reproduced their spiking activity using detailed biophysical neuronal models. Most of the putative error PCs were recorded from L3 and L5 (30/37) (See (Godlove et al. 2014) and (Sajad et al. 2019) for these methods). Compared to those recorded from L6, they showed a similar spiking profile relative to saccade onset across. Thus, we focused on modeling the activity of these two populations of neurons. We employed a model-optimization approach to estimate the excitatory pre-synaptic inputs received by these neurons around saccade onset. We described the activity of L3 error PCs using the model proposed by Eyal et al. (2018). L5 error PCs were described using the Hay et al. (2011) model, including modifications of voltage-gated calcium channel densities as in Shai et al. (2015) and hyperpolarization-activated cyclic nucleotide – HCN or Ih – channel density distribution as in Labarrera et al. (2018). The simulations included only excitatory NMDA and AMPA synaptic inputs with distributions and ratios corresponding to those in area F7d (SEF) estimated from the literature (Geyer et al. 1998; Rapan et al. 2021). The total number of NMDA and AMPA synapses per type was fixed for each simulated neuron, but their location was randomly selected in each neuron and simulation (Fig. 2A, 3A). Non-specific background pre-synaptic inputs were Poisson processes with a fixed mean (Fig. 2B, 3B).

We determined the mean of the Poisson process that replicated the observed mean inter-spike interval (ISI) distribution and spike rate of observed error PCs during the pre-target period. Inputs synchronized on saccade production were modeled as spike generators with specified temporal profiles. On a trial-by-trial basis, pre-synaptic spike times were chosen from a skewed-normal distribution (Jones et al. 2007). The number of pre-synaptic spikes and their location along the neuron, as well as their skewness, mean, and standard deviation, were optimized to reproduce the observed spiking activity of L3 and L5 error PCs.

The ISI distribution of L3 putative error PCs during the pre-target and post-saccade period followed an exponential function (Fig. 2E). In contrast, L5 putative error PCs had a uniform ISI distribution during the pre-target interval and a double exponential distribution during the post-saccade period (Fig. 3E). Additionally, L5 error PCs exhibited more bursts after errors compared to correct trials.

To estimate the background inputs to the neurons, we distinguished 3 groups of synaptic inputs according to their dendritic location—basal, oblique, and distal apical (Fig. 2A, 3A). This distinction was also used for optimizing the inputs before and after the saccade. We assumed all synapses belonging to each of these groups were activated by a Poisson process with the same mean. We simulated the spiking activity of L3 error PCs using a Poisson process with a mean equal to 3.5 for synapses located in the basal and oblique dendrites and 2.0 for synapses located in the oblique dendrites and distal apical dendrites. Fig. 2B illustrates the synaptic activation profile of a representative background input coming to the basal and oblique dendrites and the distal apical dendrites using the estimated mean of the Poisson processes. Fig. 2E shows the ISI distribution of the simulated neurons before target presentation and after the saccade for correct and error trials, respectively. The optimized biophysical models replicated the observed bursting activity (Fig. 2E). However, the minimum ISI and the normalized number of spikes count per ISI bin were larger than those observed in the experimental data. We believe this is because simulated L3 PCs could not produce ISIs smaller than 5 ms.

To reproduce the baseline activity of L5 error PCs, we only activated basal dendritic synapses with a Poisson process with a mean equal to 2 (Fig. 3B). Activation of either oblique or distal apical synapses resulted in an increase in bursting activity that was not present in the observed data during the pre-target period (Fig. 3E). While we replicated the mean spike rate of the observed L5 error PCs (Fig. 3D), the simulated ISI distributions favored ISI around 20 ms and did not show a uniform distribution (Fig. 3E). We believe the differences in the ISI distributions of observed and simulated error neurons are attributable to differences in the biophysics of the neuronal models used and variability in the biophysics of recorded neurons, not being captured by the model (see Discussion).

After optimizing the background inputs of the models, we optimized the temporal profile, location, and sequence of time-locked inputs influencing the neuron after the execution of a correct or error saccades. We calculated the mean spike rate of recorded L3 and L5 putative error PCs during the interval from 500 before to 500 ms after saccade initiation (Fig. 2D, 3D). The observed spike rate of error neurons in L3 and L5 peaked after the execution of a saccade and slowly returned to a pre-saccadic spike rate with the pre-saccadic spike rate exceeding the pre-target firing in L5 error PCs.

To evaluate the quality of the optimization, we quantified the mean amplitude, latency, and half-width of the peak (Fig. 2D, 3D). To fit these parameters, we manually optimized the time-locked inputs for the simulated neurons. The observed spiking activity of L3 error PCs was replicated by activating half of all the synapses with the same probability distribution and increasing the number of pre-synaptic spikes in error compared to correct trials (Fig. 2C). Conversely, simulated L5 error PCs were sensitive to both the location and temporal profile of pre-synaptic inputs as well as to the number of pre-synaptic spikes. Activation of oblique dendrite synapses facilitated bursting resulting in an exponential, rather than double-exponential, post-saccade ISI distribution. Thus, our final optimization of time-locked inputs only used basal and distal apical synapses (Fig. 3C).

To replicate the activity of L5 error PCs (Fig. 3D), we needed an initial basal input 70 ms before the saccade in both trial types (normal distribution: *μ* = 70 *ms*, *σ* = 140 *ms*, 2 pre-synaptic spikes) followed by a distal apical input 100 ms after the saccade (skewed normal distribution: *shape* = 2, *μ* = 100 *ms*, *σ* = 200 *ms*) and another basal input 120 ms after the saccade (right skewed normal distribution: *shape* = 5, *μ* = 120 *ms*, *σ* = 250 *ms*) (Fig. 3C). After the saccade on correct trials, all basal dendrite synapses had a probability of receiving 2 pre-synaptic spikes, and all distal apical synapses had a probability of receiving 1 pre-synaptic spike (Fig. 3C). On error trials, we needed more pre-synaptic spikes (5) coming to the second basal input around 120 ms after the saccade and a second distal apical input of 1 spike arriving 280 ms after the saccade (right skewed normal distribution: *shape* = 5, *μ* = 280 *ms*, *σ* = 250 *ms*) (Fig. 3C). The increase in sustained firing after saccades observed in L5 error PCs was produced by increasing the mean of the Poisson process for the basal background inputs from 2 to 4 after the saccade. As in the simulations of L3 error PCs, only half of the synapses received the time-locked inputs. Fig. 3E shows the observed and simulated ISI distributions of L5 error PCs in the post saccade period. In summary, we replicated the spike rate profiles of error PCs and qualitatively explained their ISI distributions during both the pre-target and post-saccade periods (Fig. 2 & 3).

### Error pyramidal cells drive midfrontal theta

Midfrontal theta is a prominent signature of performance monitoring in human EEG studies (Cavanagh and Frank 2014; Cohen 2014), being elevated on error compared to correct trials. Yet, the cellular mechanisms generating this signal are unknown. Here, we characterized the presence of theta oscillations in SEF and used biophysical modeling to ascertain whether error PCs in SEF can produce such a rhythm. First, we measured the laminar profiles of theta (θ) as well as alpha (α), beta (β), and gamma (γ) power after correct and error saccades. Because nearly all L3 and L5 putative error PCs were recorded from monkey Eu, we compared modeling results with monkey Eu’s data. Across sessions we observed a 30-74% increase in theta power after correct saccades and a 30-122% increase after error saccades (Fig. 4). This increase in theta power was significantly larger on error versus correct trials (nonparametric clustered-based permutation test, n=6, p < 0.01), extending from L3 to deep layers. Maximal θ power was observed just before the peak polarization of the error-related-negativity (Sajad et al. 2019). These results were consistent across monkeys (Supplemental Fig. S2). We also observed significantly greater α, β, and γ power on error trials in Monkey Eu but not in Monkey X (Supplemental Fig. S2). The increase was observed in the β and γ bands well after the saccade and in the α band after the saccade with a magnitude half that observed for the θ band (Supplemental Fig. S2).

**Fig. 4:**
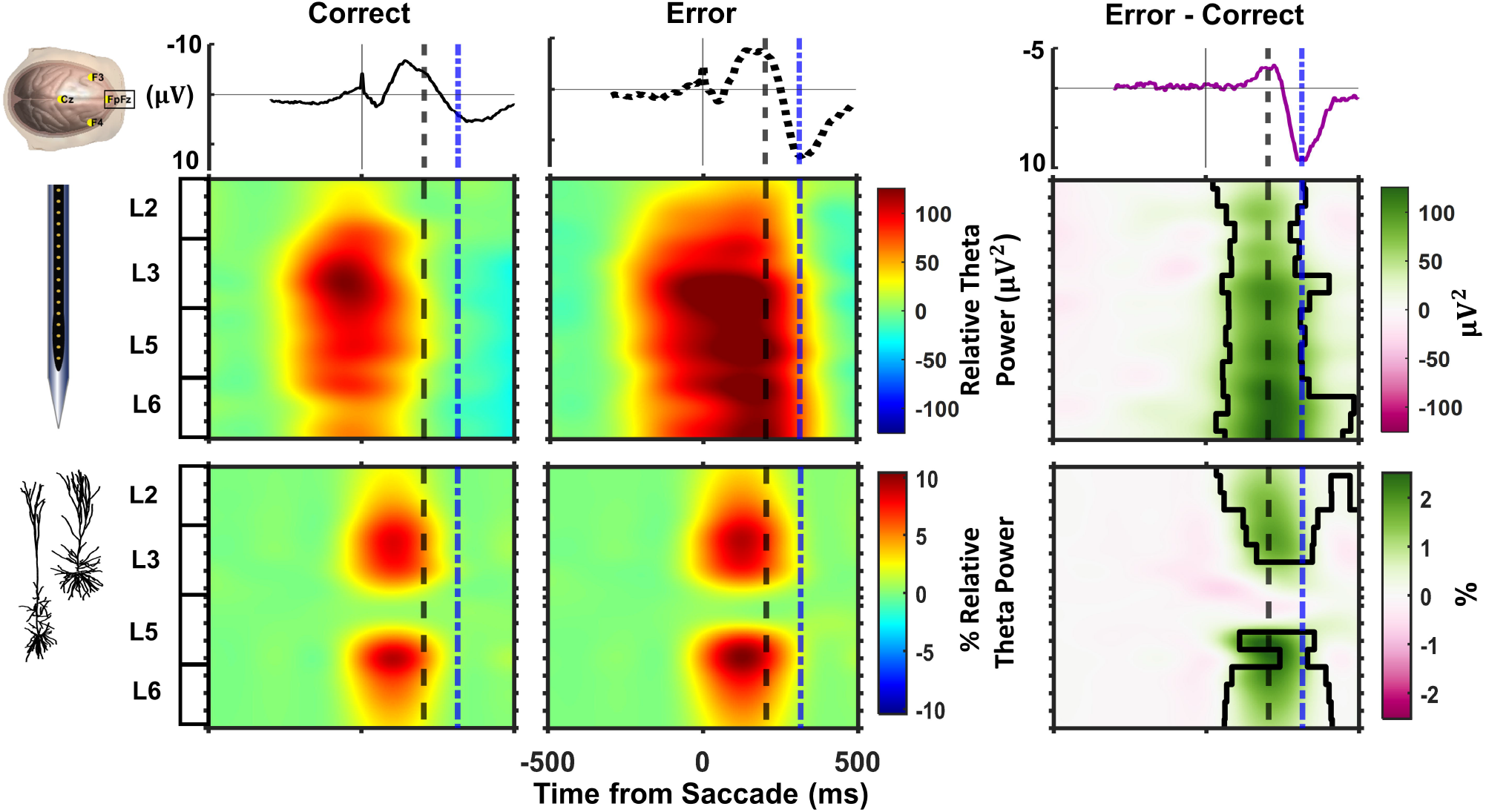
EEG and LFP θ power. Top row illustrates average ERP obtained from electrode FpFz aligned on saccade on correct (left) and error (middle) trials with the resulting difference wave (right). The spike potential associated with saccade production is evident in the correct and error plots. The difference wave highlights the ERN followed by the Pe components. The next rows plot observed (middle) and simulated (bottom) average θ power across sessions through time across the cortical layers on correct and error trials with the time-depth difference. Colormap plots power modulation relative to the mean power during 200 ms before target presentation (µV^2^) for observed and percentage of observed power. Time of peak polarization of ERN (dash) and of Pe (dot-dash) are indicated. Statistically significant regions are outlined in the difference plot.

To assess the contribution of individual error PCs to the observed increase in the laminar theta power, we simulated the activity of 625 L3 and 1,000 L5 PCs activated by random samples from the range of inputs optimized to replicate the error-related modulation and the ISI. Neuron somas were randomly positioned in L3 and L5 in a cylindrical cortical column of 3 mm diameter, with height corresponding to the vertical extent in area SEF of lower L3 (700-1100 *μm* below the pia matter) and L5 (1125-1750 *μm* below the pia matter). We calculated the LFP evoked by the activity of the simulated ensembles of error neurons at 16 equally spaced vertically aligned points in the center of the cortical column. As in the experiments, the simulated inter-electrode distance was 150μm. We compared the observed and simulated grand average laminar LFP and CSD in correct and error trials and their differences (Fig. 5, Supplemental Fig. S3).

**Fig. 5:**
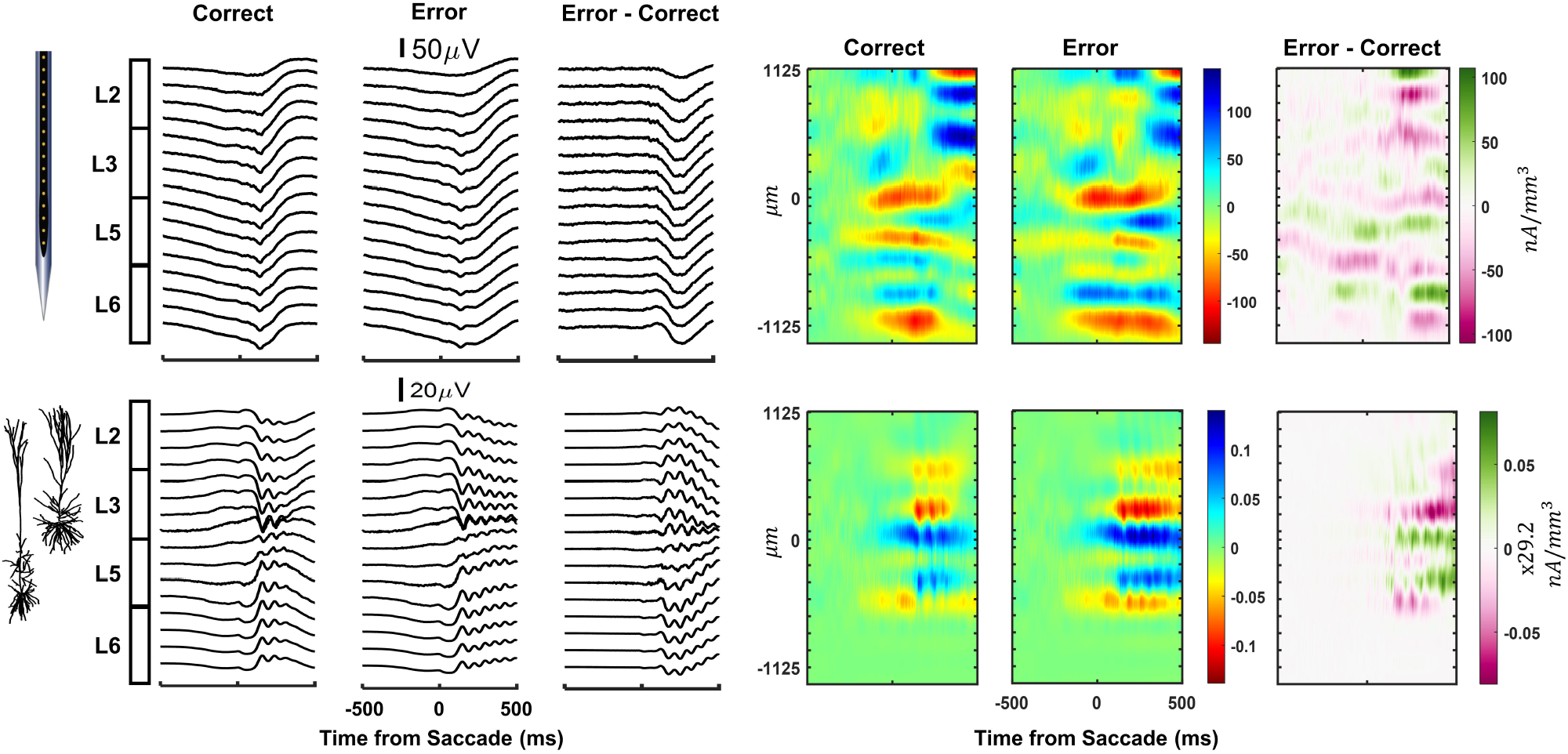
Observed (top) and simulated (bottom) average LFP and CSD. Single-trial simulated LFPs were evoked by the activity of 625 L3 and 1,000 L5 error PCs located in a cylindrical cortical column of 3 mm diameter. Neither observed nor simulated CSD had simple bipolar structure, but the simulated CSD did not replicate the observed CSD.

The observed laminar CSD could not be described as a single dipole, unlike observations in visual areas V1 (Mehta et al. 2000; Maier et al. 2011) or V4 (Herrera et al. 2022). Instead, the observed laminar CSD associated with both correct and error saccades consisted of 3 prominent sinks, one at the L3-L5 border, another in L5, and the third in deep L6. These were accompanied by a sequence of weaker, transient sinks in upper L3. Likewise, the CSD derived from the L3 and L5 error PCs simulations consisted of 3 sinks, two in L3 and one in L5. The simulated laminar CSD differed from the observed and contributed only ∼3% to the laminar CSD, even after summing CSD amplitudes over the contributes from all of the estimated total number of error neurons in these layers.

Following the same analysis pipeline, we calculated the laminar profiles of θ power relative to saccade onset on the simulated LFPs in both correct and error trials. As in the experimental data, simulated LFPs showed a significantly greater increase in post-saccadic θ power on error versus correct trials (Fig. 4; nonparametric clustered-based permutation test, n=20, p<0.01). The simulations accounted for just 10% of the observed relative laminar θ power, even without correcting by the actual number of error neurons in SEF. Analysis of the contribution of the individual populations of L3 and L5 simulated error PCs indicate that L5 error PCs, but not L3 error PCs, produce the increase in θ power around saccade. These results indicate that L5 error PCs contribute to the observed laminar θ power but, unexpectedly, contribute little to the laminar current sources observed in SEF. Additionally, the simulated L3 and L5 error PCs did not produce the laminar profile observed in the other frequency bands, suggesting these signals might be generated from other circuit mechanisms (data not shown).

To explore whether these results were associated with the intrinsic properties of the neurons, we simulated the activity of 100 unconnected L3 and L5 PCs receiving randomly activated synaptic inputs. We looked at the spectral properties of their membrane potentials and evoked LFPs in the absence and presence of the synchronized activation of all synapses 1 s after the beginning of the simulation. The voltage response of a simulated L3 and L5 PCs produced by the random input is shown in Supplemental Fig. S4. The power spectrum of the membrane potential of all L5 PCs but not L3 showed peaks in the low frequencies (θ and α bands) (Fig. 6). Inspection of the membrane potential θ phase revealed a phase-rest across L5 PCs for both dendritic and somatic membrane potential (Fig. 6B). In contrast, L3 PCs showed a phase-rest in their dendritic membrane potential, but not in their somatic membrane potential (Fig. 6A). Laminar LFP θ power maps showed an increase in θ power only for simulated L5 PCs under time-locked synchronized inputs (Fig. 6). These results indicate that L5 PCs can act as pacemakers of θ oscillations, but they are masked in the LFP unless the neurons receive a synchronized input to reset their phases.

**Fig. 6:**
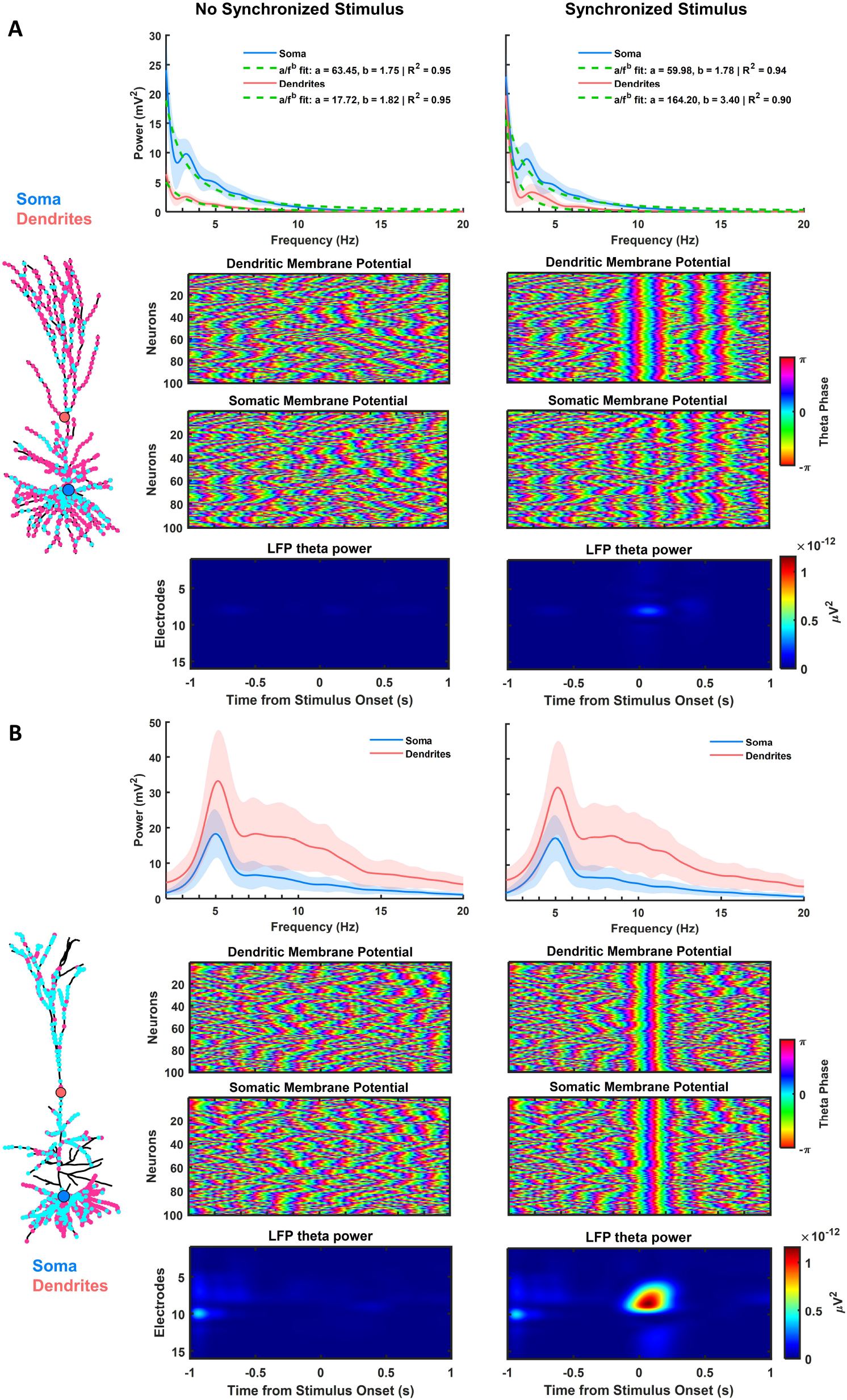
Intrinsic rhythmicity of 100 simulated L3 (A) and L5 (B) error neurons with randomized distributions of AMPA (cyan) and NMDA (pink) synapses (left) activated randomly according to Poisson processes (L3: basal mean = 2; apical mean = 1; L5: basal mean = 5, oblique mean = 4, apical mean = 1) without (left) and with (right) a synchronized input at time zero. The mean power spectra (±1.96\**SD*) of somatic (salmon) and dendritic (light blue) membrane potentials (1^st^ row) illustrate consistency of simulated neurons and pronounced peak in θ power in L5 but not L3 PCs. θ phase of the dendritic (2^nd^ row) and somatic (3^rd^ row) membrane potentials illustrate phase resetting of dendritic and soma membrane potentials of L5 neurons but only the dendritic potentials of L3 neurons. To quantify the laminar structure of θ power, the somas of the simulated neurons were randomly distributed within a cylindrical cortical column of 3 mm diameter with random depths within their associated cortical layers (L3 700-1100 μm below the pia matter; L5 1125-1750 µm). The laminar distribution of LFP θ power (bottom row) demonstrates elevated θ power derived only from L5 PCs synchronized on the phase resetting.

### Negligible contribution of error neurons to ERN current sources

Next, we employed EEG forward modeling to study SEF contributions to the ERN. We considered two current dipoles located in SEF symmetrically in each hemisphere (Fig. 7A). The temporal dynamics of the dipoles were calculated from the observed and simulated laminar CSDs (Herrera et al. 2022). We estimated EEG from the experimental CSD. Because the observed CSDs were not simple dipoles, we calculated the first three multiple moments of the observed CSDs, finding a non-zero monopole contribution arising from unbalanced current across depth. Fig. 7 shows the multipole moments obtained from the observed CSD in SEF and the multiple moments evoked by the activity of error PCs. The monopole moment of the simulated CSDs was by definition zero. In agreement with the previous results, even summing over the approximate number of error neurons in SEF, the multipole moments produced by the simulated neurons were three orders of magnitude smaller than those observed in SEF (Fig. 7). They also had different temporal dynamics (Fig. 7, Supplemental Fig. S5). Perhaps surprisingly, these results indicate a very weak and indirect biophysical contribution of SEF error neurons to the ERN. Future simulations incorporating other populations of neurons in SEF and the circuit dynamics are needed to test this hypothesis.

**Fig. 7:**
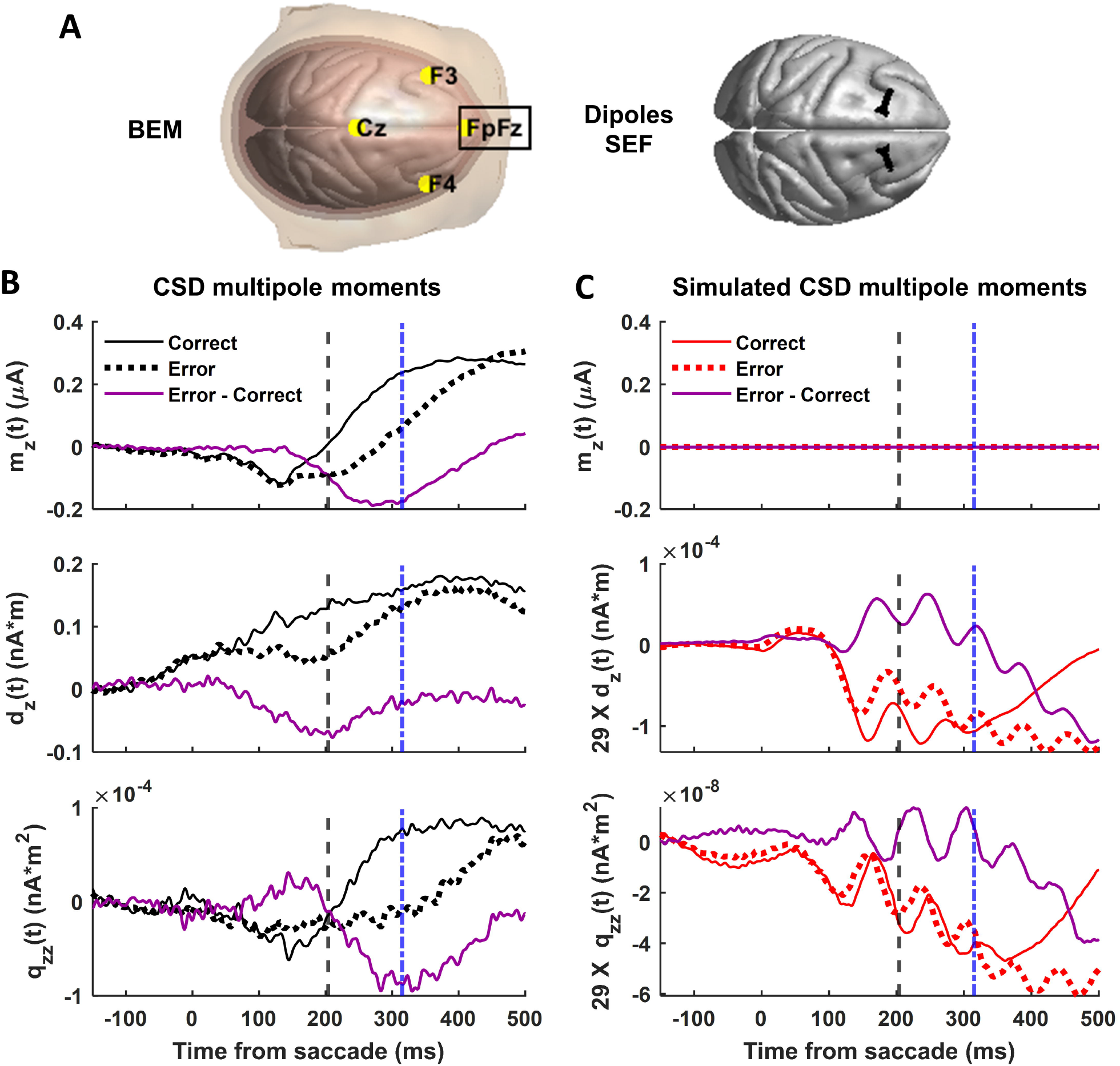
Multipole moments derived from observed (left) and simulated (right) CSD. **A.** Left – Volume conductor model of the monkey’s head (BME) with surfaces color-coded and electrodes positions in yellow. Surfaces for constructing the BEM model of the monkey’s head were obtained from the NIMH Macaque Template version 2.0 (Jung et al. 2021). right – Location of the SEF dipoles used in the EEG forward model. **B-C.** The time course of the monopole (m_z_, top), dipole (d_z_, middle), and quadrupole (q_z_, bottom) moments are plotted for correct (thin black) and error (thick dotted) trials and their difference (magenta). Time of peak polarization of ERN (dash) and of Pe (dot-dash) are indicated. The unbalanced current observed across depths creates the monopole moment. The simulated dipolar and quadrupolar moments were 4 orders of magnitude weaker than the observed. Scaling up multipoles by increasing the density of error PCs was not enough to reproduce the temporal profiles of these LFP and scalp potentials.

A complete forward model of the ERN at electrode FpFz from the SEF included the contribution of the three multipole moments (Fig. 8A, B). To account for the unbalanced currents in the observed CSD, the resulting monopole contribution was placed at the center of the equivalent current dipole in the cortical column. The monopole contribution was 1,000 times larger than the dipole and quadrupole contributions, which were of nearly equivalent magnitudes. The dynamics of the dipole moments paralleled the ERN. In contrast, the dynamics of the quadrupolar component coincided with the Pe (Fig. 8B). The SEF EEG predicted from the combination of all CSD multipole moments paralleled the dynamics of the ERN and Pe but differed substantially in magnitude (Fig. 8C). The EEG predicted by the CSD measured in SEF contributed somewhat to the ERN and weakly to the Pe component. To distinguish further the contribution of the SEF to the ERN and the Pe, we determined how the magnitude of each ERP varied with the diameter of the cortical column in which the currents were summed. As the diameter of the cortical column increased, the magnitude of the ERN varied as 1 to 4, but the magnitude of the Pe component increased as 1 to 2.

**Fig. 8:**
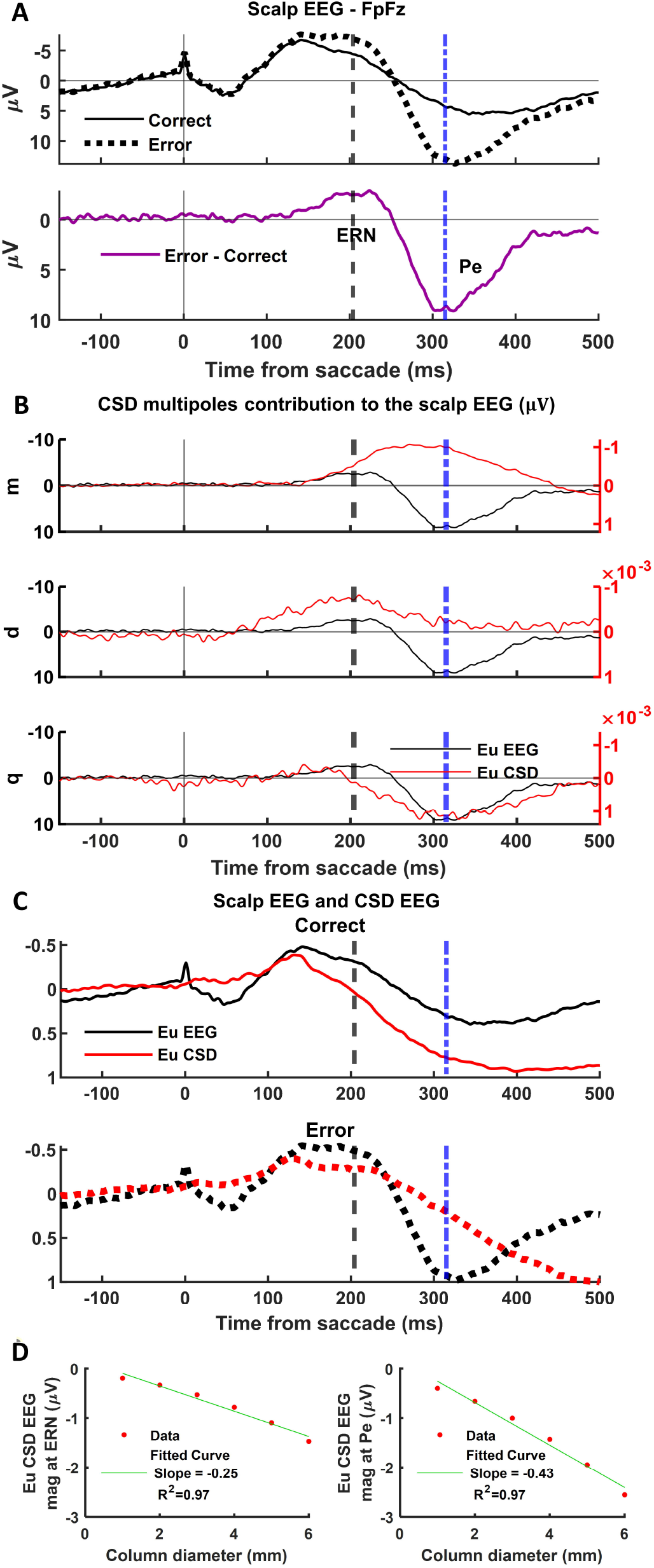
Contributions of SEF to the ERN and Pe. A. Cranial EEG during correct (thin solid) and error (thick dotted) trials with their difference (magenta) illustrating the ERN and Pe components. B. Comparison of the difference waves of the observed EEG (black, left axis) with the predicted EEG monopolar (top), dipolar (middle), and quadrupolar (bottom) components (red, right axis), respectively. The predicted EEG dipolar component explained ERN features, and the quadrupolar component reproduced those for the Pe. The presence of a monopole might indicate a more extended and diffuse neuronal activation pattern in SEF. C. Comparison of EEG observed (black) and predicted (red) from the multipolar moments derived from the CSD in SEF for correct (top) and error (middle) trials and their difference (bottom). The amplitude of the EEG signals was normalized by the maximum absolute EEG amplitude across trial types for Eu EEG and Eu CSD EEG separately. D. Variation of peak polarization of predicted ERN (left) and Pe (right) as a function of the diameter of the cortical column used in the CSD calculation. Linear regressions illustrate the significant variation, which was stronger for the Pe than the ERN.

## Discussion

The ERN and midfrontal θ have been useful biomarkers for neurological and psychiatric disorders in both basic research and clinical settings. Yet, there is a limited understanding of the cellular-level mechanisms originating these signals. Intracortical recordings in areas stipulated to be generators of these extracranial signals offer insights into their laminar origin through CSD and time-frequency analyses. However, these approaches need to be combined with detailed biophysical modeling of distinct neuronal populations to help elucidate their underlying cell-specific mechanisms. Here, we evaluated the contribution of putative error PCs to the ERN and midfrontal θ using a model-fitting approach that estimates the pre-synaptic inputs to these neurons from their spiking activity. Our biophysical model was able to capture small differences (few spikes) in the response rates of these error PCs. Our results suggest L5 putative error PCs, but not L3, contribute to error-related increases in midfrontal θ, and neither L5 nor L3 error PCs contribute biophysically to the ERN current sources. Furthermore, we estimated the SEF contribution to the scalp ERN using a multipolar expansion. Fitting well-stablished biophysical models for neocortical PCs to account for the spiking rates of error neurons with broadband spikes reinforces the conjecture that most these spikes were in fact from PCs.

### Neuron and circuit contributions to θ rhythm

Cognitive conflict detection and signaling have been associated with midfrontal θ synchronization (Cavanagh and Frank 2014; Cohen 2014). In 2014, Michael X. Cohen proposed a microcircuit model for such generation in which θ bursting, transient increase in θ power, resulted from conflict detection through L5 PCs in their apical dendrites and EEG rhythmogenesis from L3 circuital interactions between PCs and interneurons (Cohen 2014). Cohen hypothesized conflict detection mechanisms carried by L5 PCs would boost ongoing oscillations in L2/3 but would not drive such oscillations or need phase resetting by an external stimulus or response (Cohen 2014). In alignment with Cohen’s hypothesis, our model predicts a transient increase in θ power due to conflict detection in L5 PCs. However, our results show that L5 PCs’ intrinsic dynamics can drive θ oscillations and that they are only visible on the LFPs and EEG after phase-reset by synchronized synaptic inputs. These oscillations can be stronger in upper layers due to ionic channels (e.g., I_h_ and Ca^2+^ channels) present in the apical dendrites of L5 PCs.

Studies of the origin of θ oscillation in the hippocampus showed that pharmacological blockage of HCN1 (I_h_) channels or genetic deletion disrupts θ oscillations (Dickson et al. 2000; Giocomo and Hasselmo 2009; Colgin 2013; Stark et al. 2013). Additionally, it has been suggested these oscillations may originate from the inter-play between these channels and the persistent sodium (*Nap*), muscarinic *K*^+^ (*M*) and slow low threshold *K*^+^ (*K_slow_*) channels (Dickson et al. 2000; Buzsáki 2002; Wang 2010; Womelsdorf et al. 2014). The L5 PC model used in our simulations included HCN1 and M channels throughout the neuron and Nap and *K_slow_* channels only in the axon section (Leleo and Segev 2021). Hippocampal θ has also been linked to NMDA and “slow” GABA_A_ receptors (Buzsáki 2002). A recent computational modeling study found that subthalamic θ under response conflict required NMDA, but not AMPA, currents, and that the induced θ oscillations did not emerge from intrinsic network dynamics but were elicited in response to cortical inputs (Moolchand et al. 2022). Our model considered AMPA and NMDA synapses in both PC models, yet the L3 PC model did not show subthreshold membrane potential θ oscillations or induce LFP θ oscillations.

### ERN generation

The current source density derived from the simulated L3 and L5 error PCs could not explain the observed association between error-related neuron spiking and EEG of L3 but not L5 neurons (Sajad et al. 2019). Yet, error PCs contribute to the observed laminar θ power. This finding should not be surprising given the presence of neurons signaling error and reward gain/loss across SEF layers and the uncertainty about how error signals arise. One could hypothesize that error signals arrive in middle layer from thalamic afferents similar to visual afferents. However, the laminar organization of the error-related CSDs and visually evoked CSDs (Godlove et al. 2014) in SEF are not strictly dipolar as that found in sensory and visual areas. Because the error-related CSD is different from the visually evoked CSD, the error signal is not just an efferent copy arriving in middle layers of SEF.

Our EEG forward modeling indicates SEF is a neural generator of the ERN/Pe, but other sources are needed to fully explain these ERP components. The most likely source is the medial cingulate cortex (MCC), but we have little information about the laminar CSD of d/vMCC. Recently, Fu et al. (2019) found that both dMCC and pre-supplementary motor area (pre-SMA) contribute to the ERN, but at different times. Specifically, the activity of error neurons in pre-SMA preceded the activity of error neurons in dMCC. Their findings support our hypothesis that dMCC is the most likely contributor to the Pe, which could not be explained by the SEF EEG forward model. The observed CSD shows a strong monopolar component of diffuse origin, suggesting either the presence of strong electro-diffusion (Halnes et al. 2016), the existence of extended dendritic currents, or that the proposed mesoscopic source model (z-dependency) is inadequate. However, an early dipolar component explained the ERN features (i.e., peak and latency), and a late quadrupolar component peaks at Pe.

### Limitations of the model

Our model accounted for the activity of error PCs in SEF considering excitatory synaptic inputs (NMDA and AMPA), but not GABAergic inputs. We mimicked the presence of inhibitory inputs by adjusting the number and intensity of excitatory synapses. However, SEF possesses a large density of GABA receptors throughout the cortical layers (Rapan et al. 2021), which should be incorporated into the model in future studies to evaluate their role in midfrontal θ generation. In this study, we created nonparametric representations (Supplemental Fig. S6) of the laminar density of interneuron populations in SEF (i.e., calbindin “CB”, parvalbumin “PV” and calretinin “CR”). We expect the incorporation of inhibitory synaptic inputs might change the intensity and temporal profiles of the estimated inputs to the simulated error PCs, but we expect the location and timing of the inputs to be close to those estimated in this study. We also expect changes in the morphology and biophysical properties of the neuron to affect the intensity and temporal profiles of the estimated inputs.

We used neuronal models from other species (human L3 and rat L5 PCs) to reproduce the spiking profiles of the recorded neurons. Thus, we could only reproduce to some extent their ISI distributions. The L3 PC model could not fire APs with an ISI below 5ms, whereas the experimental data had a minimum ISI of 2ms. The L5 PC model produced more bursting activity than observed in the experimental data. In addition, both neuronal models produced baseline spike rates slightly larger than those for the recorded neurons. This could be associated with differences in dendritic branching between rodent/human and non-human primate neurons, resulting in different electrophysiological properties (Gilman et al. 2017; Luebke 2017; Kalmbach et al. 2018, 2021; González-Burgos et al. 2019; Galakhova et al. 2022). Additionally, our model was constrained by the data of Monkey Eu alone since Monkey X had no error neurons in L3 and L5.

In the predictions of the SEF contribution to the observed EEG, we found a current unbalance across depths in the CSD. This could be attributed to electro-diffusion given the larger density of glial cells compared to neurons in the agranular pre-frontal cortex of macaque monkeys (Dombrowski et al. 2001; Turner et al. 2016) and the presence of dendrites from nearby columns whose returning currents are not within the modeled column. As expected, we did not find a current unbalance in the simulated CSDs, which were calculated using the same methods and code as the observed CSDs. Hence, we might need to incorporate monopolar compensation in the CSD analysis to guarantee the current balance and account for such phenomena.

### Limitations of the data

We used data from two macaque monkeys recorded over three sites within SEF (one site in monkey Eu and two in monkey X). We observed differences in the laminar CSDs across monkeys, indicating a possible modular structure of SEF (Supplemental Fig. S3A). This hypothesis is supported by the presence of L3 and L5 error neurons in Monkey Eu, but not in either recording location of Monkey X. Furthermore, the laminar CSD profiles of both monkeys associated with correct and error responses were not dipolar, unlike V1, V4, and barrel cortex (Herrera et al. 2022).

### Future directions

Although our model captured the spiking profiles of L3 and L5 error PCs, future studies will benefit from the construction of neuronal models with different morphologies for macaque monkeys’ pre-frontal cortex. Additionally, sampling more sites in and around SEF will allow us to test the generality of the spiking profiles of the modeled error PCs and, hence, of our model, and evaluate the possible modular structure of SEF. Laminar recordings in d/v MCC are also needed to study its laminar organization and estimate the laminar current sources contributing to the ERN/Pe. This will allow us to formulate more complete EEG forward modeling frameworks to explain the neuronal origin of the ERN/Pe. Finally, a general CSD method that account for the existence of diffusive monopolar sources and/or compensate fictitious monopolar components must be developed to have a more reliable multiscale interpretation.

## Acknowledgments

The authors would like to thank Dr. Julio Martinez-Trujillo for useful conversations regarding the work.

## Author Contributions

Conceptualization, B.H., J.D.S. and J.J.R.; Methodology, B.H. and J.J.R.; Investigation, B.H.; Experimental design, J.D.S.; Data collection, J.D.S.; Validation, B.H.; Formal Analysis, B.H., A.S. and S.P.E.; Writing – Original Draft, B.H.; Writing – Review & Editing, B.H., A.S., S.P.E., J.D.S. and J.J.R.; Visualization, B.H.; Resources, J.D.S. and J.J.R.; Software, B.H. and J.J.R.; Funding Acquisition, B.H., J.D.S. and J.J.R; Supervision, J.D.S. and J.R.D.

## Data and Software Availability

Code for the simulations and analysis conducted in this study will be openly available on GitHub as of the date of publication. Processed data will also be available through OFS as of the date of publication. The raw data analyzed in the current study are available from Jeffrey D. Schall on reasonable request. Requests for materials should be addressed to the corresponding author Jorge J. Riera.

