## Supplemental for "Cortical Origin of Theta Error Signals"

### Supplemental Figures and Legends

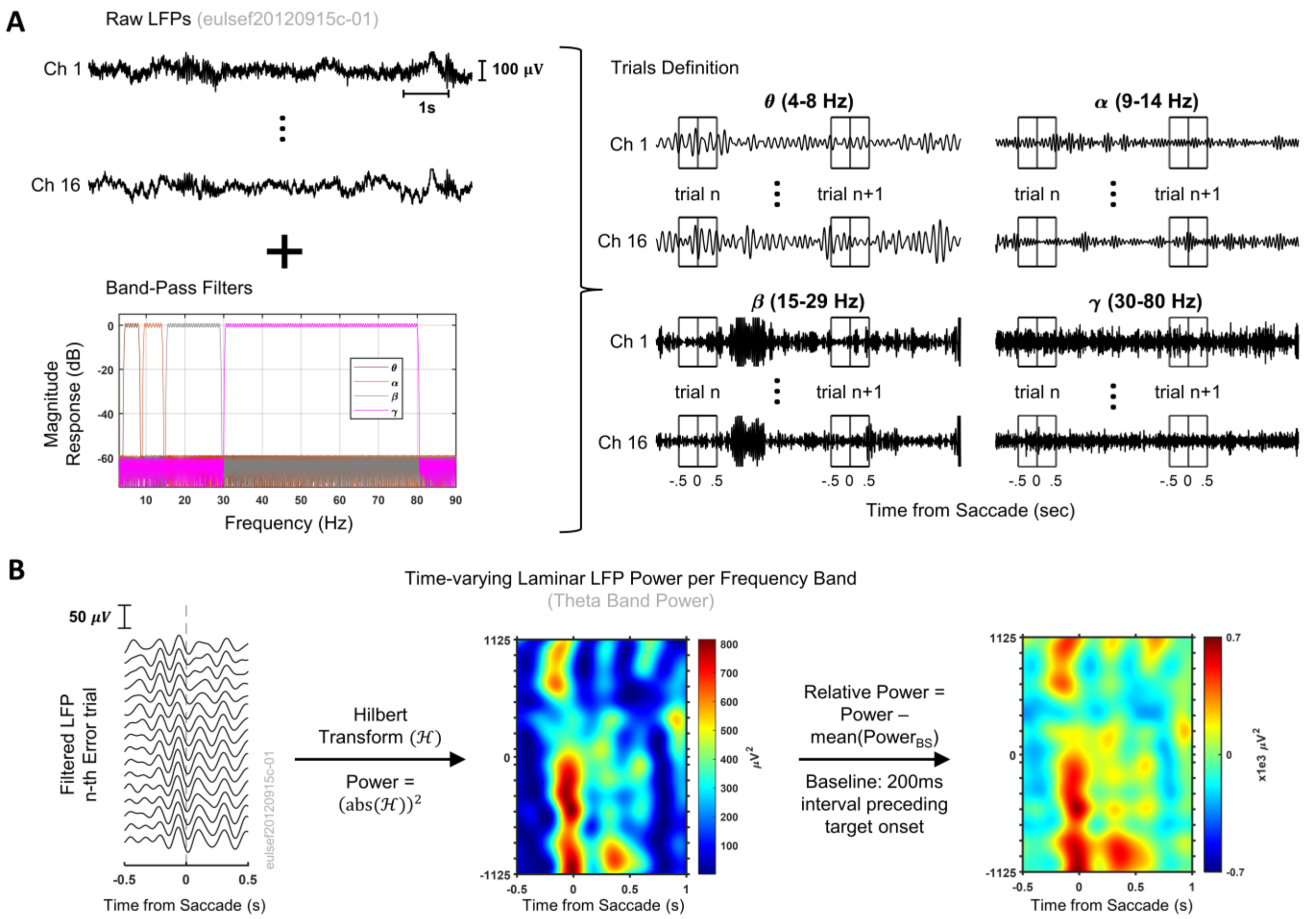


**Figure S1:** Illustration of the time-frequency analysis pipeline.

**A.** Raw LFPs (top-left) were band-pass filtered between 4-8Hz (θ band, θ), 9-14 Hz (alpha band, α), 15-29 Hz (beta band, β), and 30-80 Hz (gamma band, γ), respectively. The magnitude response of the filters is shown in the lower left panel. Filtered LFPs were epoched from -500 to 1,000 ms relative to saccade initiation (right).

**B.** The Hilbert transform of the epoched signal for each frequency band was calculated. Time-varying laminar power estimates were extracted by taking the squared magnitude of the Hilbert transform of the filtered LFPs, and baseline corrected to the mean power in the 200 ms interval preceding target onset. The final time-varying laminar LFP power maps were obtained by taking the mean across the single trial laminar power estimates. An example of the processing steps is shown for a single error trial θ-band filtered LFP.

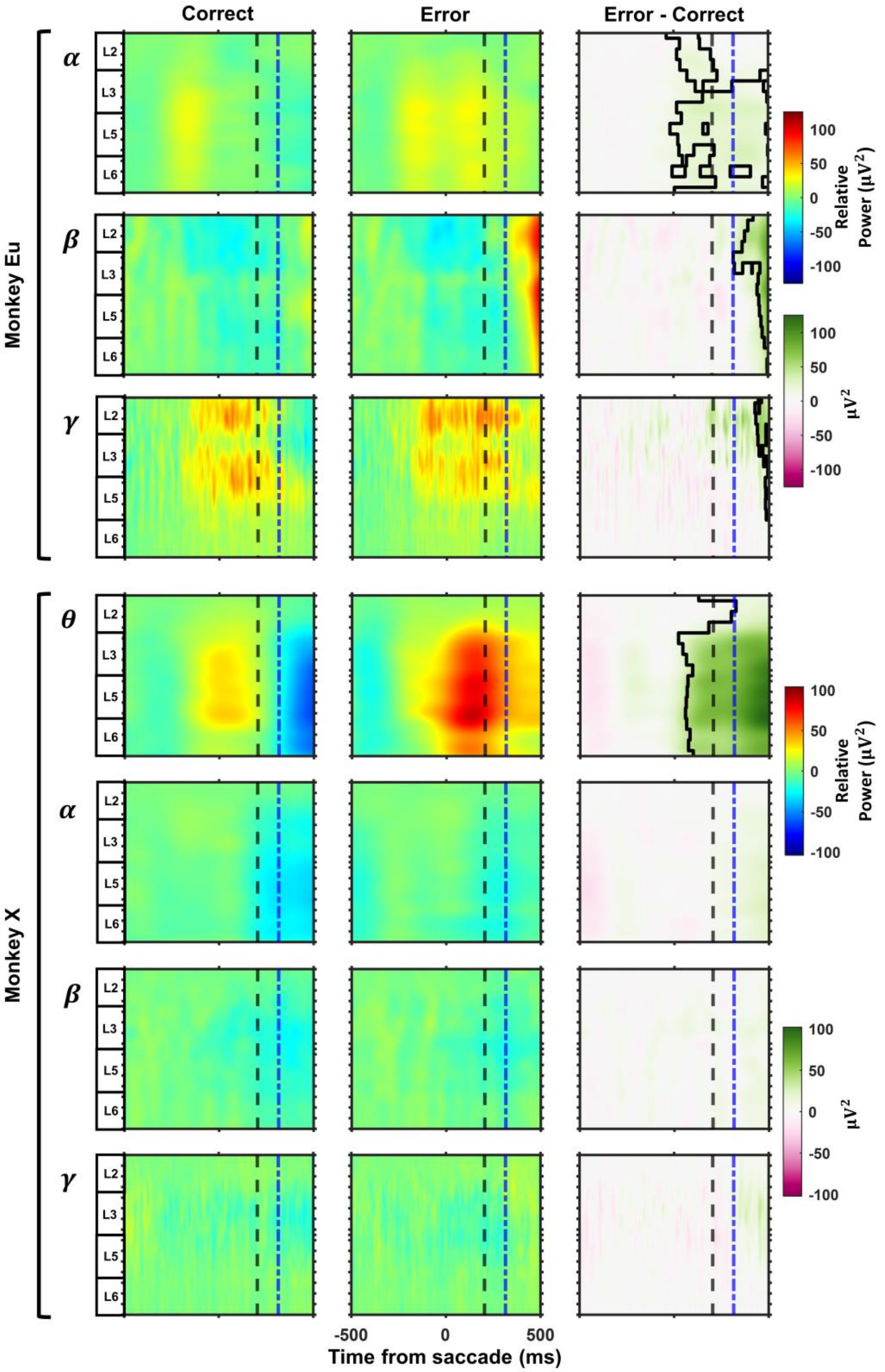

Correct
Error
Error - Correct
Monkey Eu
Monkey X
α
β
γ
θ
L2
L3
L5
L6
Relative Power ($\mu V^2$)
$\mu V^2$
100
50
0
-50
-100
-500
0
500
Time from saccade (ms)

**Figure S2:** Monkey Eu and Monkey X laminar time-varying field potential power maps per frequency band during error monitoring.

Left to right – mean LFP power map across correct trials, error trials, and their difference, respectively. Colormap represents the power modulation relative to the mean power in the baseline period (200 ms before target onset). Black traces in Error – Correct laminar power maps indicate statistically significant regions (nonparametric clustered-based permutation test). Black dashed line in all plots indicates the time of peak ERN, and blue dash-dotted line the time of peak Pe. The x-axis in all plots represents the time relative to the saccade onset, and the y-axis the cortical depth relative to the pial matter. Layers' boundaries are indicated in the left.

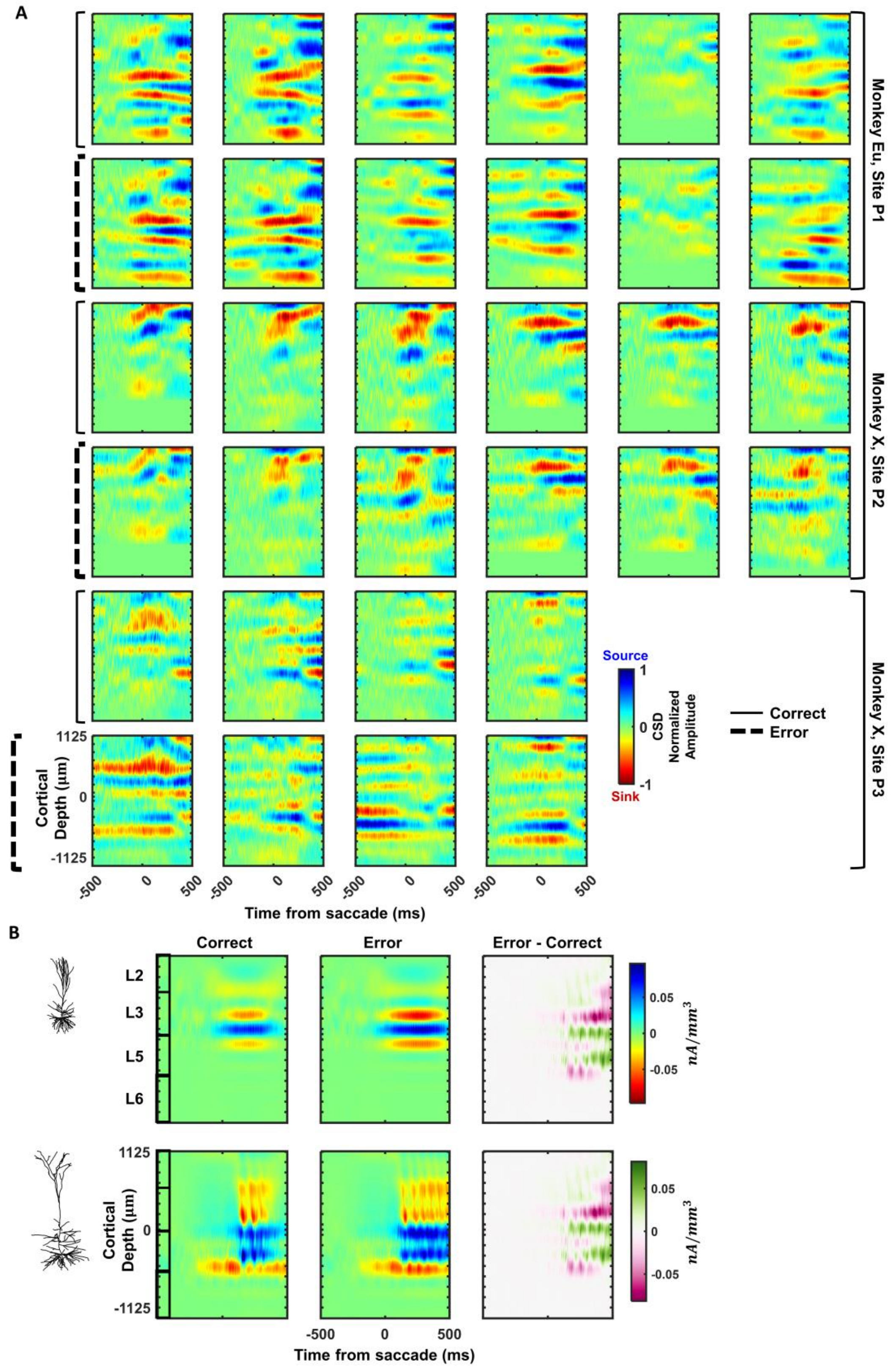
A
Monkey Eu, Site P1
Monkey X, Site P2
Monkey X, Site P3
Source
1
0
-1
Sink
CSD
Normalized
Amplitude
Correct
Error
1125
0
-1125
Cortical
Depth (μm)
-500
0
500
Time from saccade (ms)
B
Correct
Error
Error - Correct
L2
L3
L5
L6
0.05
0
-0.05
nA/mm3
1125
0
-1125
Cortical
Depth (μm)
-500
0
500
Time from saccade (ms)

**Figure S3:** Laminar Current Source Density Maps.

**A.** Single sessions laminar current source density maps. Thick dotted lines on the left indicate Error trials and thin solid lines Correct trials. The monkey and recording location to which each session belongs to are indicated on the right. The x-axis represents the time relative to saccade onset. CSD amplitudes were normalized by the maximum absolute amplitude across trial types per session. The y-axis in all plots represents the cortical depth relative to the L3/L5 boundary, depth-zero.

**B.** Individual contributions of the activity of L3 (top) and L5 (bottom) error pyramidal cells populations to the simulated CSDs. Layers' boundaries are indicated on the left. In all panels, blue denotes current sources and red current sinks.

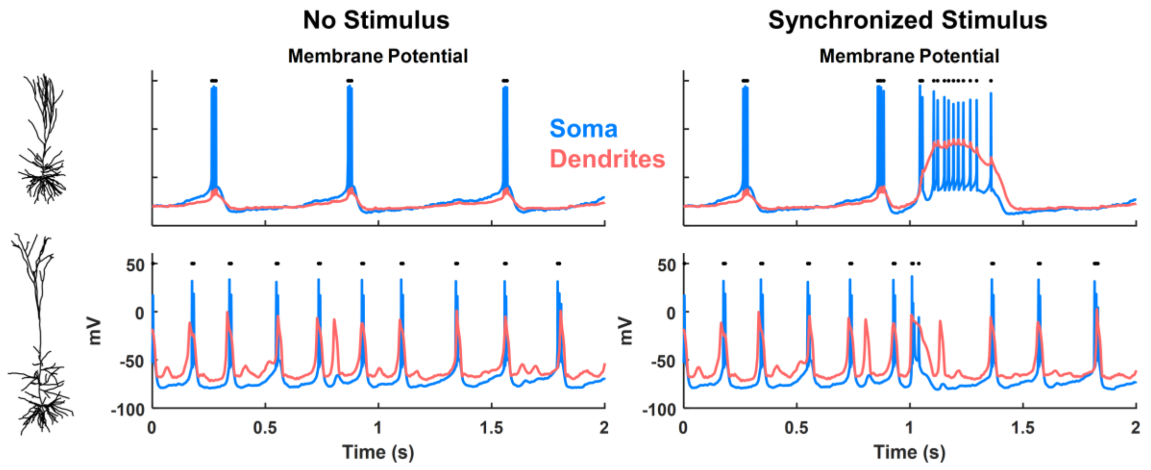


**Figure S4:** Voltage response of a simulated L3 and L5 pyramidal cells (in Fig. 6) to excitatory synaptic inputs randomly activated by a Poisson process (L3 PCs – mean equal 2 and 1 for basal and apical synapses, respectively; L5 PCs – mean 5, 4, and 1 for the basal, oblique, and distal apical synapses, respectively) without (left) and with (right) a synchronized stimulus at 1s.

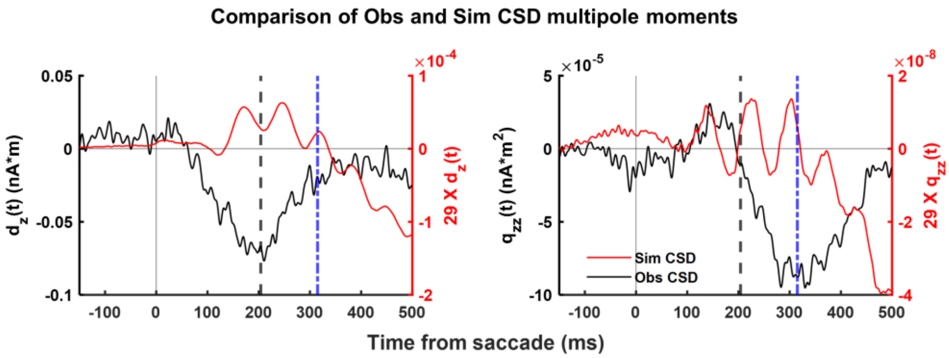


**Figure S5:** Comparison of the experimental (black) and simulated (red) CSD dipole (left) and quadrupole (right) moments. Black dashed line in all plots indicates the time of peak ERN, and blue dash-dotted line the time of peak Pe. Lines represent the difference between Error and Correct trials for each multiple moment.

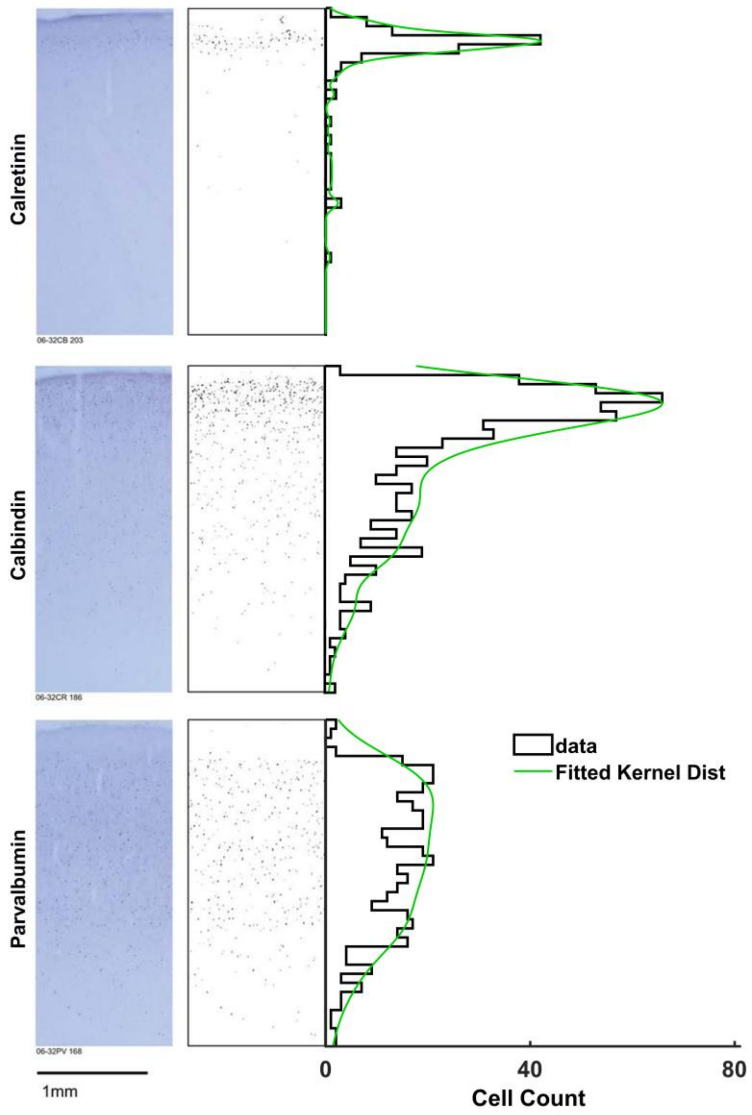


**Figure S6:** Laminar distribution of GABAergic interneurons in Supplementary Eye Field (SEF). Related to Discussion.

Left – Coronal sections through SEF immunohistochemically reacted for the calcium binding proteins Calretinin – CR, Calbindin – CB, and Parvalbumin – PV.

Right – Location of positive neurons (dots) throughout SEF cortical laminae with associated histogram of cell counts and estimated laminar distributions (green lines). Cells were identified though a semi-automatic classification routine (see Godlove et al. (Godlove et al. 2014) for details on the algorithm). Laminar distributions were estimated in MATLAB using the function fitdist() and considering a Kernel probability density function ($\widehat{f_p}(z, h_p) = \frac{1}{n_p h_p} \sum_{i=1}^{n_p} K\left(\frac{z - z_i}{h_p}\right)$ where $p = \{CR, CB, PV\}$; $z_1, \ldots, z_{n_p}$ represent the depth of each cell in the *p-th* population, $n_p$ is the sample size or total number of cells in the *p-th* population throughout the cortical laminae, $K(z) = \frac{1}{\sigma\sqrt{2\pi}} e^{-\frac{(z-\mu)^2}{2\sigma^2}}$ is the kernel smoothing function ($\mu = 0$, $\sigma = 1$), and $h > 0$ is the bandwidth or width of the kernel). The estimated bandwidth for CR, CB, and PV interneurons was $h_{CR} = 0.0494$, $h_{CB} = 0.1428$, and $h_{PV} = 0.2327$, respectively.

The figure was adapted from Fig. 9 of Godlove et al. (2014) published in the Journal of Neuroscience under a Creative Commons Attribution-Noncommercial-Share Alike 3.0 Unported License (CC-BY-NC-SA). Histology images and plots of the location of positive neurons were not modified. New histograms of cell count were created for estimating the laminar distributions of CR, CB, and PV interneurons.
